# Barcoded overexpression screens in gut Bacteroidales identify genes with new roles in carbon utilization and stress resistance

**DOI:** 10.1101/2022.10.10.511384

**Authors:** Yolanda Y. Huang, Morgan N. Price, Allison Hung, Omree Gal-Oz, Davian Ho, Héloïse Carion, Adam M. Deutschbauer, Adam P. Arkin

## Abstract

A mechanistic understanding of host-microbe interactions in the gut microbiome is hindered by poorly annotated bacterial genomes. While functional genomics can generate large gene-to-phenotype datasets to accelerate functional discovery, their applications to study gut anaerobes have been limited. For instance, most gain-of-function screens of gut-derived genes have been performed in *Escherichia coli* and assayed in a small number of conditions. To address these challenges, we developed Barcoded Overexpression BActerial shotgun library sequencing (Boba-seq). We demonstrate the power of this approach by assaying genes from diverse gut Bacteroidales overexpressed in *Bacteroides thetaiotaomicron*. From hundreds of experiments, we identified new functions and novel phenotypes for 29 genes involved in carbohydrate metabolism or tolerance to antibiotics or bile salts. Highlights include the discovery of a D-glucosamine kinase, a raffinose transporter, and several routes that increase tolerance to bile salts through lipid biosynthesis. This approach can be readily applied to develop screens in other strains and additional phenotypic assay types.

## Introduction

The past decade has seen massive sequencing initiatives and -omics studies on the human gut microbiome.^1–4^ We now appreciate that there is tremendous complexity in the network of interactions between host and microbes, and among microbes in this ecosystem. Despite their importance for human health, we lack a detailed mechanistic understanding of microbial functions in the gut.^5, 6^ The vast majority of bacterial genes have not been experimentally characterized, which in combination with their high functional diversity, has led to a knowledge gap in the field.^7, 8^ Accurately predicting gene functions often requires both a mechanistic understanding of the protein family and supporting phenotypic data, which can be difficult to obtain.

Functional genomic libraries can rapidly connect genotypes to phenotypes. Moreover, DNA barcoding of individual strains allows for parallel analysis of strain fitness in genome-wide libraries through barcode sequencing (BarSeq).^9^ In recent years, randomly barcoded transposon sequencing (RB-TnSeq) mutant libraries have enabled high-throughput genetic screens across hundreds of conditions in a cost-effective manner and have accelerated functional assignments of genes in diverse bacteria, including the human gut anaerobe *Bacteroides thetaiotaomicron* (*B. theta*).^10–12^ Gain-of-function screens are complementary to loss-of-function screens and have the advantage of capturing genes from bacteria that have yet to be cultivated or genetically modified. However, most libraries expressing gut-derived DNA have relied on *E. coli* as the overexpression strain, which is phylogenetically divergent from members of the abundant gut phyla Bacteroidota (formerly Bacteroidetes) and Bacillota (formerly Firmicutes).^13–18^ In particular, promoter recognition of *Bacteroides* genes in *E. coli* can be limiting and *Bacteroides* encode unique ribosomal binding sites (RBSs), which can present barriers for protein expression in *E. coli*.^19–21^ Therefore, Bacteroidales serve as more suitable overexpression platforms to study functions in the gut, yet few genetic screens have been performed in them.^22, 23^

Another challenge in gain-of-function screens is the reliance on deep sequencing or isolation of individual strains to identify beneficial genes.^13–18, 22^ This is time-consuming and laborious to perform for hundreds of assays, which has limited these approaches to a small number of conditions. Random DNA barcoding of libraries can drastically increase the scale of overexpression screens. For example, dual barcoding of vectors (Dub-seq) has been used to construct libraries expressed in *E. coli*, but these are currently limited to ∼250,000 strains and are restricted to DNA sources with sequence assemblies, which are required for mapping.^24–27^

Here, we exploit advances in long-read sequencing to develop a new workflow termed Boba-seq, or <u>B</u>arcoded <u>O</u>verexpression <u>BA</u>cterial shotgun library <u>seq</u>uencing, to perform large-scale gain-of-function screens using DNA barcoded vector libraries. We demonstrate this with overexpression in *B. theta*, although this approach is readily applicable to alternative expression strains and is agnostic to the source of DNA being assayed. Bacteroidales are major constituents of the gut microbiota that establish stable, long-term associations with the human host. They play crucial ecological roles in the gut, including metabolism of diverse, host- recalcitrant carbohydrates and modulation of the human immune system.^28^

We constructed Boba-seq libraries from seven Bacteroidales and first validated our method in a complementation screen. We then assayed a pool of six libraries in competitive fitness assays and identified genes involved in the metabolism of 15 carbon substrates and tolerance to seven inhibitory compounds (antibiotics or bile salts). Genes with novel functions or phenotypes included enzymes, transporters, and regulators. In addition to gene function discovery, our work enables large-scale gain-of-function screens in a clade of gut bacteria important for human health.

## Results

### Overview of workflow to build and assay barcoded overexpression libraries

The Boba-seq workflow utilizes a single DNA barcode per plasmid to increase the throughput of phenotypic assays. This approach consists of five general steps (Fig. 1). First, we construct replicative vectors to encode an inducible promoter, an insertion site for DNA of interest, a terminator, a DNA barcoding site, and an origin of replication for the overexpression host. Empty vectors are barcoded by cloning in random 20 nucleotide barcodes.^29^ In the second step, randomly sheared and size-selected DNA fragments from the organism(s) of interest are cloned into the barcoded vectors. Here, fragment sizes can be optimized to capture single genes or larger gene clusters that might encode the target phenotypes. Next, fragments are linked to unique barcodes using long-read sequencing. This simplifies the mapping step to a single PCR amplification step instead of the complex TnSeq-like protocol that was used previously to map dual-barcoded vectors.^24^ In step four, libraries are conjugated into the overexpression strain. In the final step, libraries are screened across a panel of conditions where the inoculum (Time0) and samples grown in selective media are harvested for BarSeq. Inserts that lead to increased strain fitness are identified by calculating fitness scores for each barcode, which represents the log2 change in the relative abundance of the strain. Fitness scores are then filtered to identify potential causative genes for the observed fitness gain.

**Figure 1.**
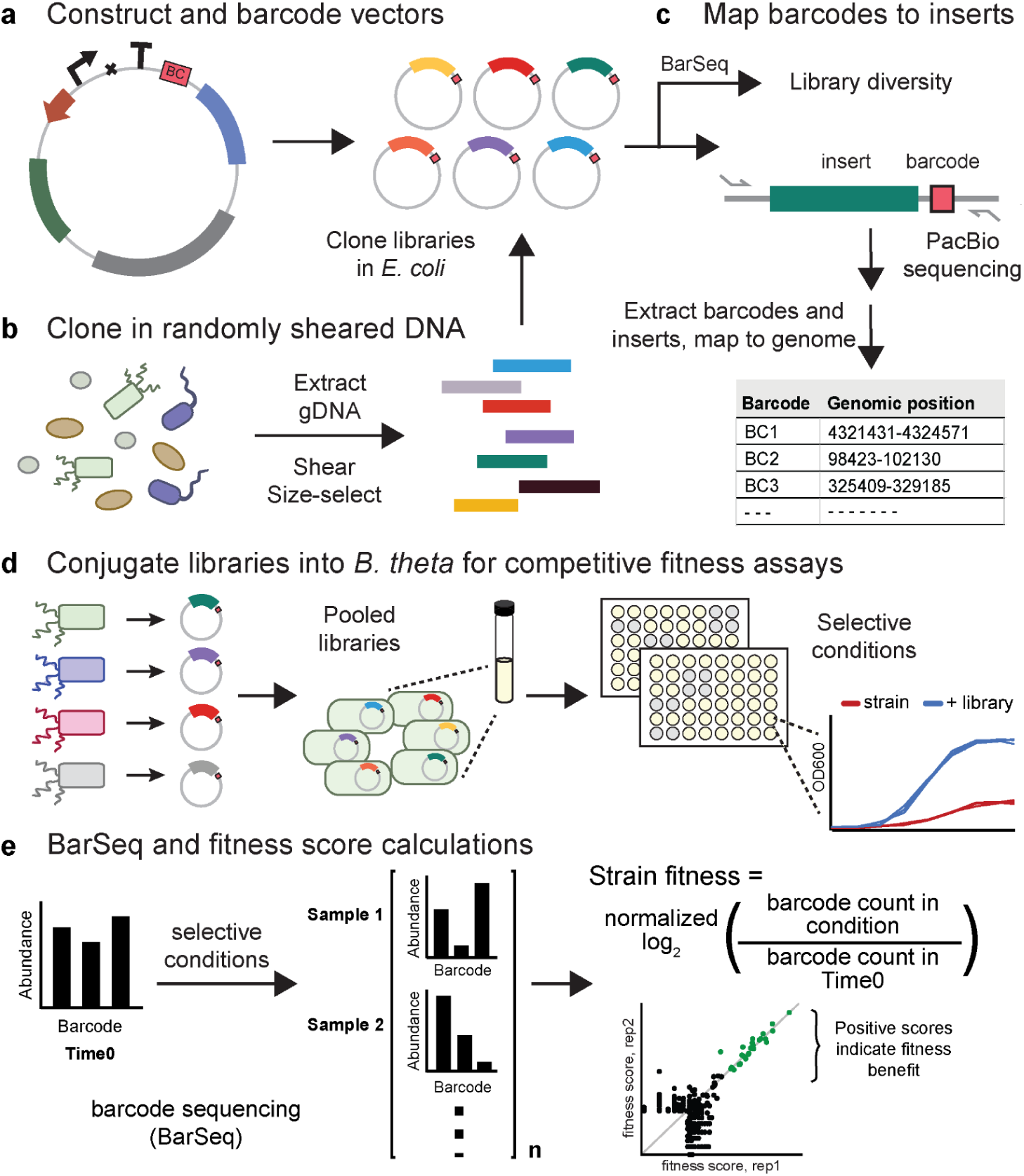
The Boba-seq workflow for high-throughput functional screens. A) Clone DNA barcodes into replicative vectors for the overexpression host. B) Extract DNA, shear, size- select, and clone into barcoded vectors. C) Quantify library diversity using BarSeq. Map barcodes to inserts using long-read sequencing technology and generate a barcode-to-insert mapping table. D) Conjugate mapped libraries into the overexpression strain for fitness screens. Libraries can be pooled to increase the genetic diversity assayed. A range of selective conditions can be used to determine fitness differences across strains, such as carbon utilization and stress tolerance. E) Quantify library composition before and after growth on selective conditions using BarSeq. A fitness score is calculated for each barcode or strain. Significantly high fitness scores in both replicates point to putative beneficial genes in the condition.

### Construction of DNA barcoded overexpression libraries from Bacteroidales genomic DNA

We generated and characterized new shuttle vectors for *B. theta* VPI-5482 (see Methods).^30–33^ This led to three replicative vectors: pNBU2_repA1, pNBU2_repA2, and pNBU2_repA3 (Fig. 2a, Extended Data Table 1). Our vectors contain an anhydrotetracycline (aTc) inducible promoter, a ribosomal binding site (RBS), and a terminator.^33–35^ Inducible gene expression by aTc was verified using NanoLuc reporter protein (Extended Data Fig. 1a). The plasmid copy number for these vectors in *B. theta* was determined to be around 3, 7, and 8, respectively, using droplet digital PCR (Extended Data Fig. 1c). Collectively, these vectors are replicative in 10 of the 13 Bacteroidales tested, including *B. theta* (Extended Data Fig. 1d). The vector pNBU2_repA1 was then barcoded to contain approximately 14 million unique barcodes (see Methods). Insert lengths of ∼3 kb were selected to capture at least one gene on most fragments. In total, seven libraries were generated using gDNA from *B. theta*, *B. caccae*, *B. fragilis*, *B. salyersiae*, *B. uniformis*, *Parabacteroides johnsonii*, and *Parabacteroides merdae*, which share a maximum of 92% average nucleotide identity. Phylogenetic relationships between these seven strains are shown in Fig. 2b. 81% of all protein-coding genes in these genomes have at least one protein homolog (≥40% a.a. identity, ≥75% coverage), either in the same genome or in another genome.

**Figure 2.**
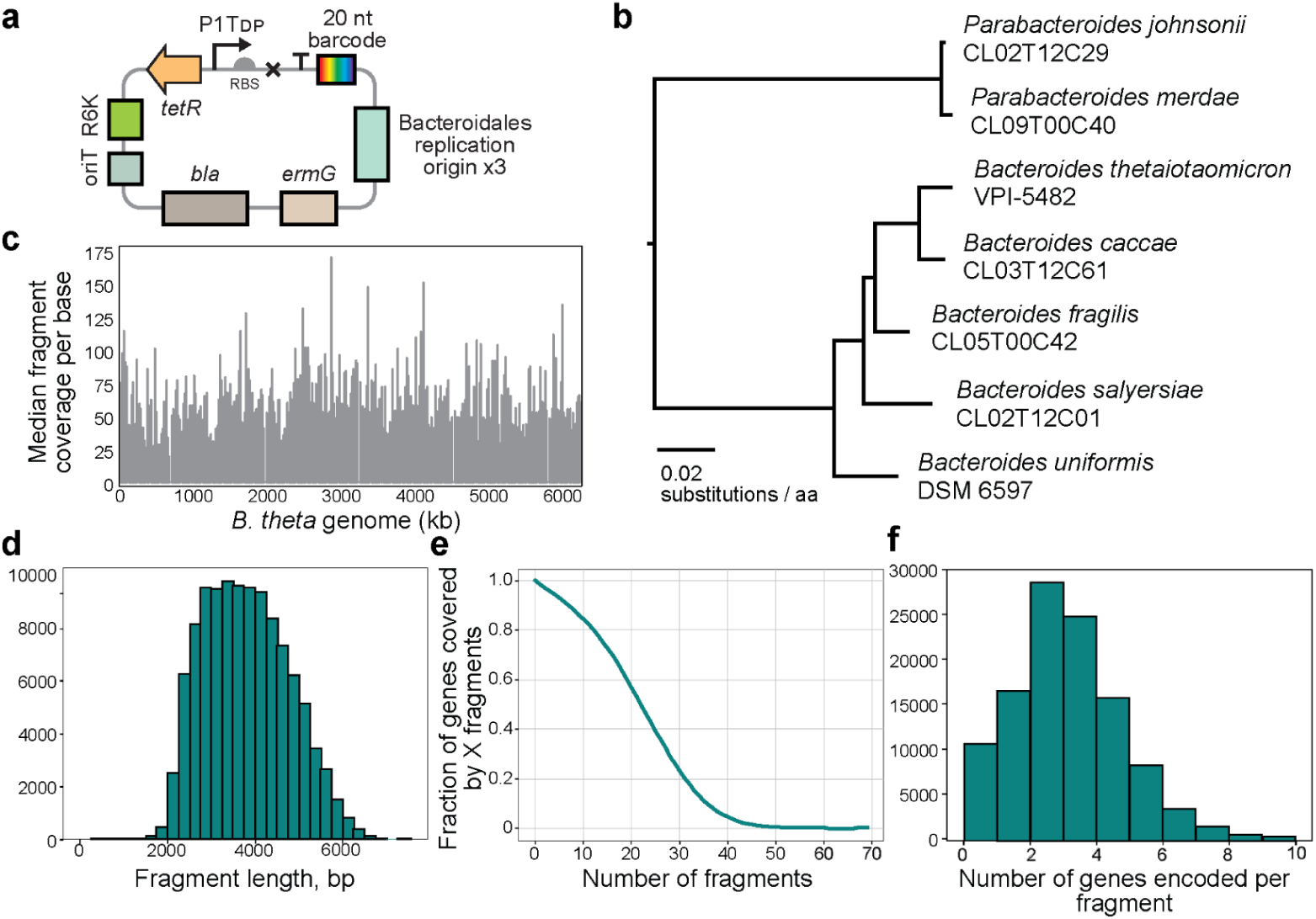
Boba-seq libraries were constructed from gDNA of *Bacteroides* and *Parabacteroides* isolates with high genomic coverage. A) Three replicative vectors were constructed by modifying an integrative vector (pNBU2-ErmG).^33^ B) 24 universal bacterial marker genes from the Genome Taxonomy Database (GTDB) were used to generate this phylogenetic tree to show phylogenetic relatedness of the seven isolates used to build Boba-seq libraries.^36^ Protein sequences of marker genes were aligned with HMMer 3.3.1 (hmmer.org) and then a tree was inferred with FastTree 2.1.11^37^ and midpoint rooted. Coverage of the *B. theta* genomic library: C) fragment coverage per base across the genome (NC_004663.1) plotted in 20 kb windows, D) histogram of fragment lengths, E) cumulative gene coverage plot, and F) histogram of number of full-length genes encoded per fragment.

Next, we developed a pipeline to link barcodes to fragments on each vector (Extended Data Fig. 2). We performed PacBio amplicon sequencing the insert-barcode region to obtain 10–24-fold sequencing depths (Extended Data Table 2). 77–91% of barcodes detected by BarSeq were mapped, hence only a small fraction of each library is not analyzed. On a per genome basis, fragments cover 87–98% of protein-coding genes with 76–477 protein-coding genes unmapped (Extended Data Table 2, Fig. 2c). In all libraries, the missing proteins are generally longer than mapped proteins (median 1,758 vs. 927 bp) and 5% have a nearly-identical copy (≥95% a.a. identity) elsewhere in the genome. Overall, 50% of the missing proteins are over 2 kb in length or are duplicated, which reflects limitations imposed by fragment sizes and difficulties in mapping highly similar regions. After excluding genes of over 1.5 kb or with duplicates, 31% of the remaining genes have a homolog in the same or another library that is also missing (Extended Data Table 3). Conserved missing genes suggest possible toxicity in *E. coli* and include proteins that are resistant to horizontal gene transfer such as ribosomal subunits, although the majority cannot be readily explained based on the annotation.^38^

Libraries consist of barcode diversities ranging from 40,911–97,471 and median fragment lengths of 2.7–3.7 kb (Fig. 2d, Extended Data Fig. 3a, Extended Data Table 2). The majority of fragments encode 1–2 complete genes and the median fragment coverage per gene is 12–24 (Fig. 2e,f, Extended Data Fig. 3b,c, Extended Data Table 2). Overall, 50,000 fragments of ∼3.1 kb are sufficient to cover a bacterial genome of approximately 5–6.5 Mbp with a median fragment coverage of >10. A sequencing depth of >10✕ with long reads can map a majority of barcodes. In summary, we constructed seven libraries with high genome-wide coverages.

**Figure 3.**
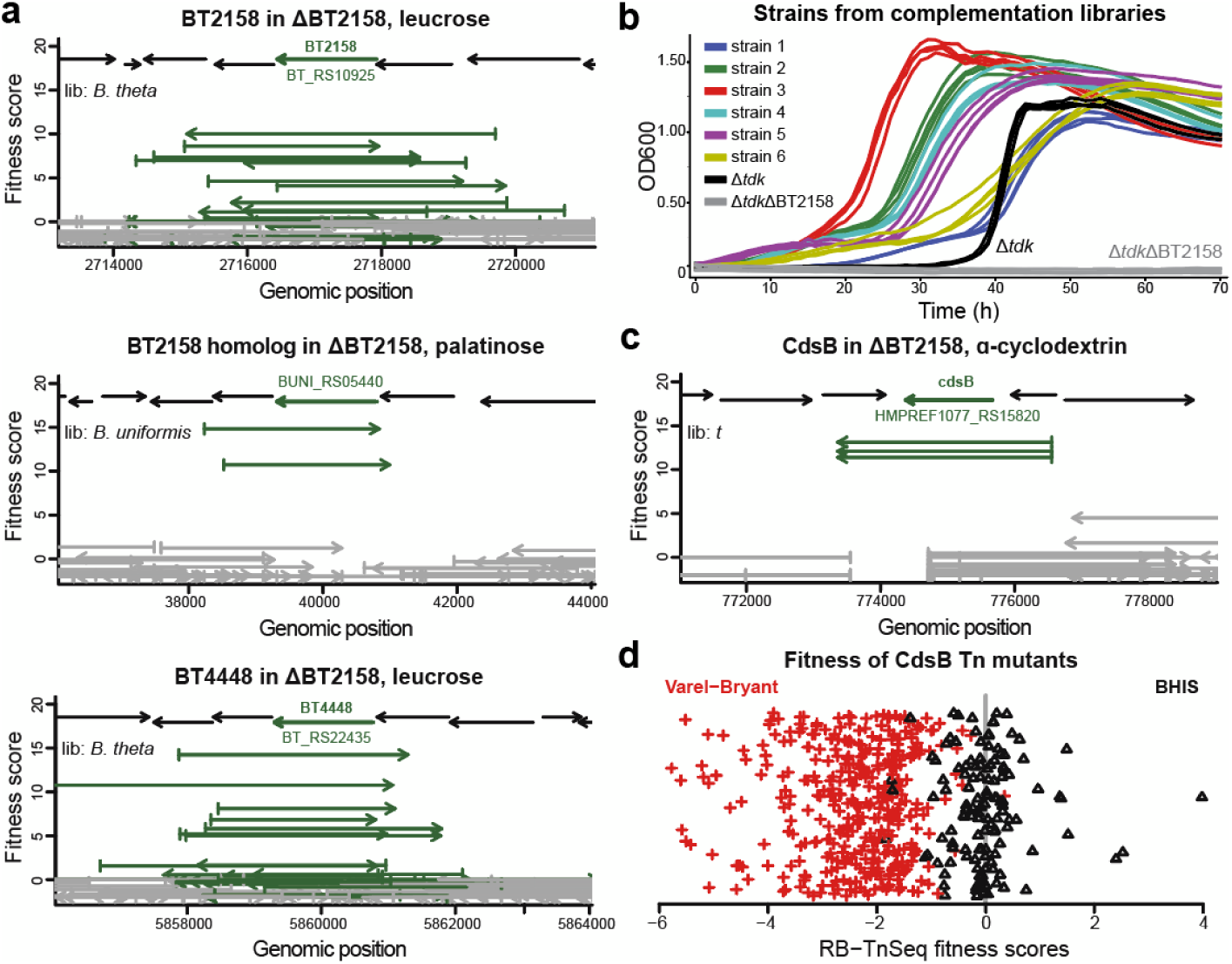
Fitness data from complementation assays. BT2158, its homologs, and cysteine degrading enzymes were enriched in complementation assays. The average fitness score from both replicates is plotted for each fragment. Fragments that cover the complete gene are indicated in green. A) BT2158 and BT4448 in *B. theta*, and a homolog (BACUNI_RS05440), were beneficial for the ΔBT2158 mutant during growth in leucrose and palatinose. B) Growth curves of ΔBT2158 mutant complemented with either BT2158 or a homologous hit. Complemented isolates, the Δ*tdk* parental strain, and the Δ*tdk*ΔBT2158 mutant were cultured in the VB medium with 20 mM trehalose. Strains 1–2 encode BT2158, strains 3–4 encode BT4448, and strains 5–6 encode BACUNI_RS05440. See Extended Data Table 6 for the precise genomic boundaries of cloned fragments. Each strain was grown in replicates of 4. OD600 values are pathlength-corrected and blank-normalized. C) CdsB was enriched in the ΔBT3703 mutant in ɑ-cyclodextrin. D) RB-TnSeq fitness scores for CdsB (BT2080/BT_RS10540) are averages across all 37 CdsB transposon mutants in the *B. theta* library and are log2 fold-changes that were previously reported (x-axis).^11^ Scores are highlighted in red for assays in the VB minimal medium and in black for the BHIS rich medium. The y-axis is random scatter for visualization.

### Complementation assays as a proof-of-concept

To validate our workflow in a phenotypic screen, we complemented *B. theta* ΔBT2158 and ΔBT3703 with *B. theta, B. fragilis, B. uniformis*, and *P. johnsonii* libraries.^11^ BT2158 (BT_RS10925) encodes a periplasmic glycoside 3-dehydrogenase required for disaccharide and glucosinolate metabolism.^11, 39^ ΔBT2158 does not grow on trehalose (glucose-α-1,1- glucose), leucrose (glucose-α-1,5-fructose), or palatinose (glucose-α-1,6-fructose) (Extended Data Fig. 4). The second mutant lacks the ɑ-glucosidase SusB (BT3703/BT_RS18670) part of the starch utilization system and has reduced growth on ɑ-cyclodextrin compared to wild-type.^11^ We then inoculated each library in a defined minimal medium (Varel-Bryant or VB) containing a single carbon substrate in the presence of the inducer aTc (Extended Data Table 4).^40^ Among these four source genomes, three encode BT2158 or a homolog, and three encode BT3703 or a homolog, which are expected to provide fitness gain in the conditions tested.

We computed fitness scores for each fragment using the log2 fold-change of each barcode’s relative abundance between the selective condition and the Time0. We focus on high positive scores because Boba-seq is a gain-of-function assay and we established filters to identify biologically consistent hits. First, fragments with a fitness score ≥5 and a z score ≥4 in both replicates are considered statistically significant. Using these thresholds, no significant scores were found in a control comparison of Time0 samples (see Methods). Second, we consider each hit to be biologically consistent if it is confirmed by overlap or by protein similarity. Specifically, a hit is biologically consistent if 1) there is an overlapping fragment with significant high scores or 2) there is a protein homolog (≥40% identity) from a different genomic region (from within the same library or a different library) with a significant fitness score. Finally, we inspect the gene(s) covered by these fragments to identify the causative gene. This led to 16 biologically consistent region by condition pairs encoding 15 unique proteins that conferred a fitness gain in leucrose, palatinose, or ɑ-cyclodextrin (Extended Data Table 5).

For libraries expressed in ΔBT2158, growth in leucrose or palatinose led to the expected hits encoding BT2158, its nearly-identical paralog BT4448 (BT_RS22435), or a homolog (BACUNI_RS05440) from *B. uniformis* (Fig. 3a). In trehalose, we observed extremely high fitness scores (>12) for strains encoding each of these genes, but each strain had high fitness in just one replicate, so they did not meet our criteria for statistical significance. These strains likely grew quickly early on in one replicate, but were outcompeted by other fit strains in the second replicate. To verify our results, we isolated clones encoding either BT2158, BT4448, or BACUNI_RS05440 and observed improved growth on trehalose (Fig. 3b). Of note, three of six fragments encode genes oriented opposite to the inducible promoter with <50 bp upstream and were likely transcribed from a weak, unexpected promoter located downstream of the vector insert site (Extended Data Table 6, Extended Data Fig. 1b).

Previously, it was observed that the ΔBT2158 mutant cannot grow on these disaccharides, suggesting that BT2158 is essential for metabolism and BT4448 cannot rescue its activity.^11^ In contrast, our data show that BT4448 can functionally complement ΔBT2158 when expressed from a vector. BT4448 is likely not expressed to sufficient levels in these conditions to rescue growth in ΔBT2158 and that overexpression from our vector led to higher activity.

Complemented ΔBT3703 libraries in ɑ-cyclodextrin led to a cysteine desulfidase (CdsB) and a cysteine β-lyase as beneficial hits (Fig. 3c, Extended Data Table 5). Here, the selection for cysteine degrading enzymes appears to be stronger than for BT3703 and its homologs since ΔBT3703 is capable of growth on ɑ-cyclodextrin.^11^ Cysteine degradation is likely beneficial due to toxicity of the 8.25 mM cysteine in the minimal medium (Extended Data Fig. 5).^41, 42^ This is also supported by RB-TnSeq data for *B. theta*, where disruption of CdsB (BT2080/BT_RS10540) led to reduced fitness across all conditions in the same minimal medium (Fig. 3d).^11^ Similarly, cysteine β-lyase mutants of *B. theta* exhibited mild fitness defects in many conditions.^11^ Of note, the rich medium used for *B. theta* RB-TnSeq fitness assays also contains 8.25 mM cysteine, and thus this effect is specific to minimal media conditions (Fig. 3d).^11^ Cysteine is commonly added to Bacteroidales growth media as a reductant, but our result cautions against its use at a high concentration in minimal media.

In summary, we established a set of stringent filters to identify beneficial hits. In particular, complementation worked well for conditions with strong selective pressure where the host strain exhibits little or no growth.

### Fitness profiling of pooled libraries using BarSeq

After establishing a pipeline for fitness data analysis, we sought to map genotypes to phenotypes in conditions relevant for the gut habitat using wild-type *B. theta* as an expression host. In total, we performed 222 assays consisting of 42 single carbon sources in a defined minimal medium and 16 inhibitory compounds at 2–5 concentrations in a rich medium (Extended Data Table 4). The vast majority of assays were carried out in the presence of the inducer aTc. Carbon sources included dietary and host-derived complex polysaccharides, gelatin, mucins, and simple carbohydrates. Inhibitory compounds consisted of antibiotics and bile salts. Six libraries (excluding the *B. theta* library) were pooled into a single inoculum (or Time0). Growth was observed in 29 carbon substrates and all stress conditions. 192 samples were processed for BarSeq and included two replicates per condition and two Time0 samples per fitness screen.

Of the 305,382 uniquely-mapped barcodes present in the six libraries in the conjugation donor strain, we detected 283,816 barcodes in our fitness assays (having at least 2 reads across all assays and controls). Across all libraries, 21,042 protein-coding genes were assayed. We identified 1,172 barcodes with statistically significant scores using the same criteria as previously detailed for the complementation screen (Fig. 4a). We identified biologically consistent fragments for 120 unique regions across 22 compounds. The majority of beneficial regions were curated to assign a putative causative protein or proteins by inspecting fitness score vs. fragment coverage plots (Extended Data Fig. 6). Experimental evidence from any characterized homolog was assessed using PaperBLAST and taken into consideration as well.^43^ Overall, hits across all conditions consist of 76 protein clusters with 30 clusters containing multiple homologs at ≥40% amino acid identity (Extended Data Table 5).

**Figure 4.**
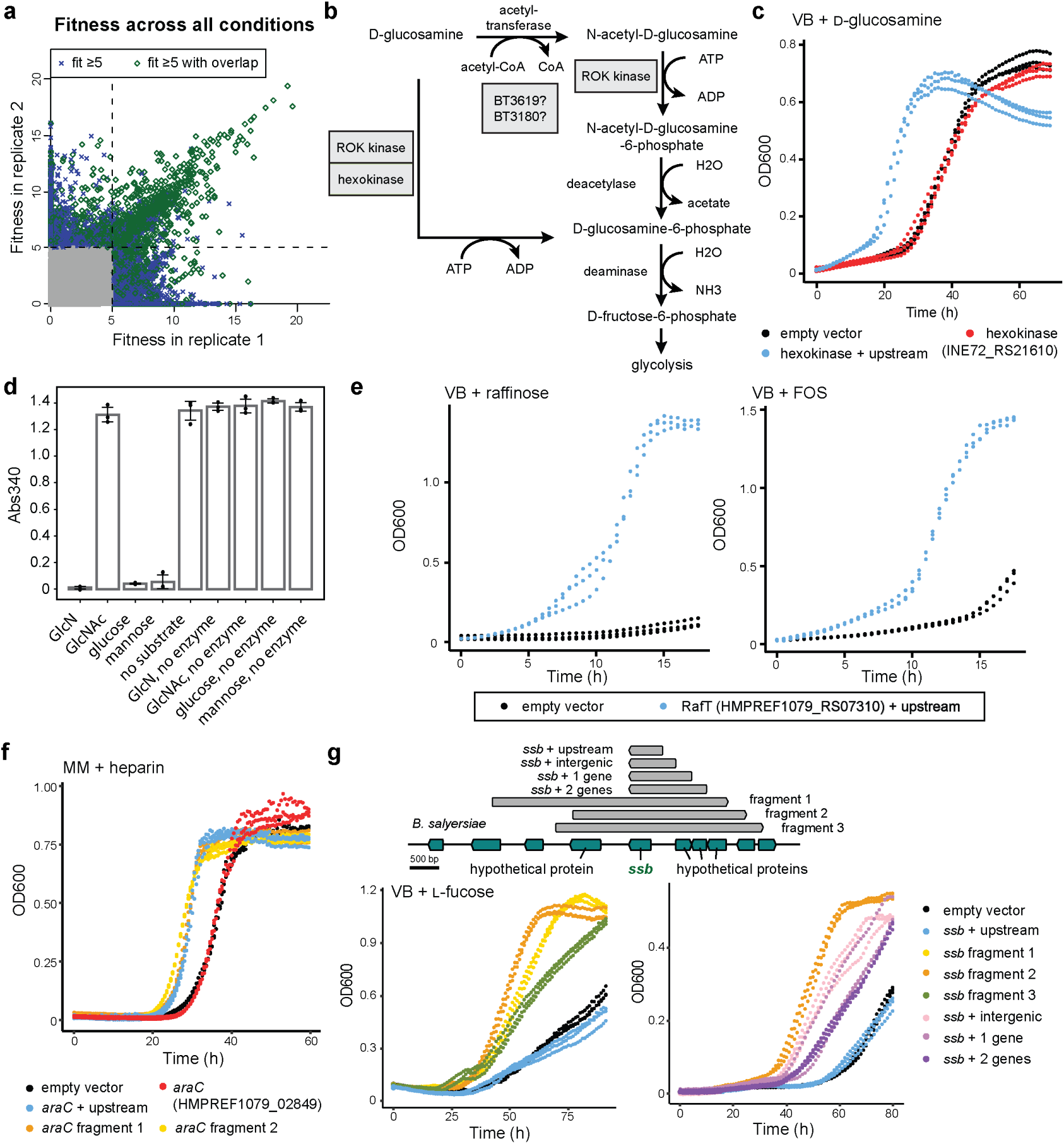
Competitive fitness assays in wild-type *B. theta* uncovered novel functions in carbon utilization. A) Consistency of strain fitness values between two replicates across 93 pairs of experiments with 6 pooled libraries. Grey points represent fitness scores <5. Dashed lines separate fitness scores of ≥5. Barcodes with fitness scores ≥5 whose regions overlap with other high-scoring barcodes are highlighted in green. B) D-Glucosamine and GlcNAc share a metabolic pathway in Bacteroidales. ROK kinase and hexokinase hits from our screen are displayed based on biochemical evidence. C) A hexokinase conferred a growth advantage on D-glucosamine when overexpressed in *B. theta*. D) *In vitro* end-point enzyme assays coupled to NADH depletion were performed for hexokinase to reveal a substrate preference for D-glucosamine (GlcN), glucose, and mannose. E) *B. theta* expressing RafT is more fit growing on raffinose and FOS as the sole carbon source. F) An AraC regulator provides a fitness gain on heparin. G) A genomic region containing a ssDNA-binding protein (HMPREF1071_RS19060) is beneficial on L-fucose. For all growth curves, individual strains were cloned to identify the causative region and constructs with “+ upstream” indicate the inclusion of the upstream 200 bp. The VB minimal medium was used for each condition except for heparin where a different defined minimal medium (MM)^44^ was used (Extended Data Table 4). See Extended Data Table 6 for the genomic boundaries of library fragments verified by growth assays of individual strains.

Consistent with the complementation screen, we detected high fitness scores for CdsB and cysteine β-lyase across 9 conditions containing cysteine (Extended Data Table 5). In addition, 21 carbon substrates were tested using an alternative reductant, dithiothreitol (DTT). While most of these assays led to similar hits, after excluding Cys-degrading enzymes, assays with DTT generally led to more hits compared to assays with cysteine.

We noticed that a large number of strains did not show a benefit even though they encode a curated causative gene or genes (Extended Data Table 5). Specifically, 31% of strains that contain the putative causative gene(s) show a significant benefit (fitness score ≥5 and z ≥4 in ≥1 replicate). This is likely partly due to differences in gene expression caused by variations in fragment boundaries: for strains with the exact same insert sequence where at least one strain has a statistically significant score, 70% have a significant score in at least one replicate. This is much higher than the 31% rate for all strains with causative genes. Our vector’s inducible promoter was designed to ameliorate expression issues, but many genes are likely transcribed from their native promoters, which are not always included in fragments. Moreover, most protein-coding genes in Bacteroidales do not use a Shine-Dalgarno sequence for translation initiation, therefore the vector’s promoter or RBS could interfere with translation.^21, 45^

### High-confidence hits uncover metabolic enzymes in carbohydrate utilization

Here, we focus on metabolic enzymes beneficial on single carbon sources. Three substrates led to expected hits based on known functions or annotations (Extended Data Table 5). These include the L-fucose catabolic operon from L-fucose and a β-glucosidase from D-cellobiose and D-salicin (Extended Data Fig. 6).^11, 46^ We verified that the β-glucosidase confers a growth benefit when expressed in *B. theta* on D-cellobiose and D-salicin (Extended Data Fig. 7a).

Additionally, an ɑ-glucosidase (INE72_RS06280) in the glycosyl hydrolase family 31 was beneficial in palatinose in both the pooled library assays and when overexpressed in individual strains (Extended Data Fig. 6, 7b). Since palatinose contains an α-1,6-glycosidic bond, we annotate this enzyme as an α-1,6-glucosidase.

An ROK family (repressor, open reading frame, kinase) protein and a hexokinase were both important for growth on D-glucosamine. We first show that strains encoding the ROK kinase or the hexokinase have a growth advantage in D-glucosamine (Fig. 4c, Extended Data Fig. 7c). In support of our data, a ROK kinase homolog in *B. theta* (31% identity) can phosphorylate D-glucosamine and N-acetyl- D-glucosamine (GlcNAc) *in vitro*.^47^

In *B. theta*, D-glucosamine and GlcNAc share a metabolic pathway where D-glucosamine can be acetylated to form GlcNAc (Fig. 4b).^48^ GlcNAc is then phosphorylated and deacetylated to form D-glucosamine-6-phosphate.^49^ While two kinases were beneficial for growth on D-glucosamine, no hits were observed in GlcNAc, which may be a result of lower selective pressure since *B. theta* grows better on GlcNAc than D-glucosamine in the VB medium. The hexokinase is important for glucose and mannose metabolism as previously reported using a *B. fragilis* deletion mutant (AY664812, 79% identity).^50^ While our fitness data support D-glucosamine as an additional substrate, we cannot exclude the possibility of GlcNAc phosphorylation by this hexokinase since it is an intermediate of the metabolic pathway (Fig. 4b). Therefore, we purified the hexokinase (INE72_RS21610) to establish its activity in a coupled enzyme assay. The hexokinase has a strong substrate preference for glucose, mannose, and D-glucosamine, with little or no activity toward GlcNAc (Fig. 4d). Here, we use both genetics and biochemical approaches to identify a new D-glucosamine kinase and demonstrate its importance in D-glucosamine metabolism in Bacteroidales.

### TonB-dependent transporters mediate raffinose uptake

The TonB-dependent transporter (TBDT) family can carry out active transport of vitamins, siderophores, peptides, and glycans across the outer membrane, and are prevalent in the phylum Bacteroidota, with 121 putative TBDTs in *B. theta* alone.^51^ Two TBDTs were uncovered to be beneficial for growth in raffinose and fructooligosaccharides (FOS) (Extended Data Table 5, Extended Data Fig. 6). To our knowledge, this is the first report of raffinose transporters in Bacteroidales. Raffinose is a trisaccharide part of the raffinose family of oligosaccharides (RFOs) that is widespread in dietary plants and abundant in the standard mouse chow used in microbiome studies.^52, 53^ A distantly-related TBDT from *B. theta* (BT1763/BT_RS08935, ≤28% identity) is involved in the uptake of FOS with β-2,6 linkages, but the FOS substrate used in our study contains predominantly β-2,1-linkages (Extended Data Table 4).^54^

We name these transporters RafT as they share similarity at 49% identity (HMPREF1077_RS18310, HMPREF1079_RS07310), but have distinct gene neighborhoods. An α-galactosidase is present downstream of *Pj-*RafT (from *P. johnsonii*) and can likely release galactose from raffinose.^55^ A β-D-fructofuranosidase is genomically adjacent to *Bf-*RafT (from *B. fragilis*) and can hydrolyze raffinose.^56^ Both nearby enzymes are predicted to be periplasmic, which supports a direct uptake of raffinose by the RafTs. This is also supported by RB-TnSeq data for a RafT homolog in *Phocaeicola vulgatus* (HMPREF1058_RS00615, >47% identity) that is important for raffinose utilization (unpublished data). In addition to raffinose, *Bf*-RafT was also a hit in FOS and in the complementation screen for leucrose. Leucrose, raffinose, and FOS all contain D-fructose as a terminal residue, which might mediate substrate recognition by *Bf-*RafT. We confirmed that *Bf-*RafT is beneficial for *B. theta* grown on raffinose and FOS (Fig. 4e). Collectively, the combination of data from Boba-seq libraries, RB-TnSeq libraries, gene neighborhoods, and growth experiments, support raffinose uptake as a function of these TBDTs.

### A regulator confers fitness in growth on heparin

In addition to genes involved in metabolic and transport pathways, our results point to regulatory mechanisms as well. An AraC family transcriptional regulator (HMPREF1079_RS07695) conferred a fitness benefit on heparin in both pooled fitness assays and individual strain growth assays (Fig. 4f, Extended Data Fig. 6). This protein has been associated with resistance to DNA damaging agents, but a role in heparin utilization or more broadly, carbohydrate metabolism, had not been reported.^57, 58^

### A mobile genetic element contributes to competitive growth on L-fucose

L-Fucose is a key nutrient for Bacteroidales in the gut with sources including dietary polysaccharides and host glycoproteins such as mucins. In addition to being a carbon source, L-fucose is a component of bacterial capsular polysaccharides and fucosylated glycoproteins.^59^ During nutrient-limiting conditions, *B. theta* can induce fucosylation of host glycans to increase nutrient availability for resident microbes, thus altering gut microbiota composition.^60, 61^ Given the importance of this molecule for host-microbe interactions, we sought to identify genes that provide a competitive growth advantage on L-fucose.

Here, L-fucose catabolic genes were beneficial across all L-fucose conditions (Extended Data Table 5). Unexpectedly, a genomic region encoding a ssDNA-binding protein (SSB) was also detected among all L-fucose conditions (Extended Data Fig. 8a). Additionally, there are three hypothetical proteins upstream of *ssb* in this conserved region, which is part of a ∼50 kb putative mobile genetic element (MGE) present in Bacteroidales (Extended Data Fig. 8b). In support of this, a conserved recombinase and resolvase are encoded nearby. First, we verified our fitness data using three high-scoring fragments and observed growth benefits on L-fucose (Fig. 4g). Then, by using truncated fragments, *ssb* and the upstream intergenic region appears sufficient for a partial fitness gain (Fig. 4g).

SSB proteins play important roles in DNA replication, recombination, and repair, and have multiple protein binding partners.^62^ Additionally, they are present on plasmids and play anti-defense roles by inhibiting SOS response and by protecting transferred ssDNA from nuclease-catalyzed degradation.^63^ Based on known functions of SSB, we could not readily propose a mechanism to account for the observed growth advantage on L-fucose. We also cannot exclude the possibility that the upstream intergenic region might contribute to the phenotype. Therefore, in an attempt to gain molecular insights, we performed an RNA-seq experiment using the most fit strain encoding a library fragment and a strain with an empty vector (Extended Data Fig. 8c, Extended Data Table 7, *p*-value <0.05, Wald test). Transcripts for L-fucose catabolic genes are not differentially abundant between these two strains, which rules out a direct regulatory mechanism by which this region is conferring a benefit.

A partial restriction endonuclease subunit S (BT4522/BT_RS22805) was strongly upregulated in the strain expressing the *ssb* fragment (log2 fold-change or log2FC of 5.5). This is caused by phase-variation in the Type I restriction modification system and is likely catalyzed by the adjacent recombinase. Based on transcript read alignments, the partial S subunit gene recombined with an upstream S subunit gene such that it became expressed as part of the new S subunit (Extended Data Fig. 8d). We also observed expression differences in capsular polysaccharide (CPS) biosynthesis, with the CPS3 locus upregulated (average log2FC of 1.46) and the CPS5 locus repressed (average log2FC of −4.6). Both CPS loci are regulated by invertible promoters in addition to the upxY transcriptional anti-terminator and upxZ *trans* locus inhibitor system.^64^ Consistent with its known role, we speculate that the SSB affects recombination events and altered phase-variation at these loci. However, we cannot pinpoint a direct mechanism by which these transcriptional changes might impart a benefit on L-fucose. Despite this, we uncovered a novel link between this conserved region within a MGE and a growth advantage on L-fucose.

### Multiple strategies confer resistance to antibiotics

The gut microbiota is under dynamic selection pressure from host-secreted factors, dietary compounds, and xenobiotics. Therefore, we examined genes important for fitness in the presence of different classes of antibiotics, bile salts, and an antioxidant (Extended Data Table 4). Seven inhibitory compounds led to significant hits with greater fitness gains typically in the highest concentrations (Extended Data Table 5).

First, we identified three known genes (*cepA*, *bexA*, *folA*) important for antibiotic resistance through antibiotic inactivation, drug efflux, or increased target expression (Fig. 5). Additionally, a pentapeptide repeat-containing protein hit on ciprofloxacin is similar to a quinolone-resistance protein (∼30% identity, EF0905) that inhibits quinolone binding to DNA gyrase and topoisomerase.^65^

**Figure 5.**
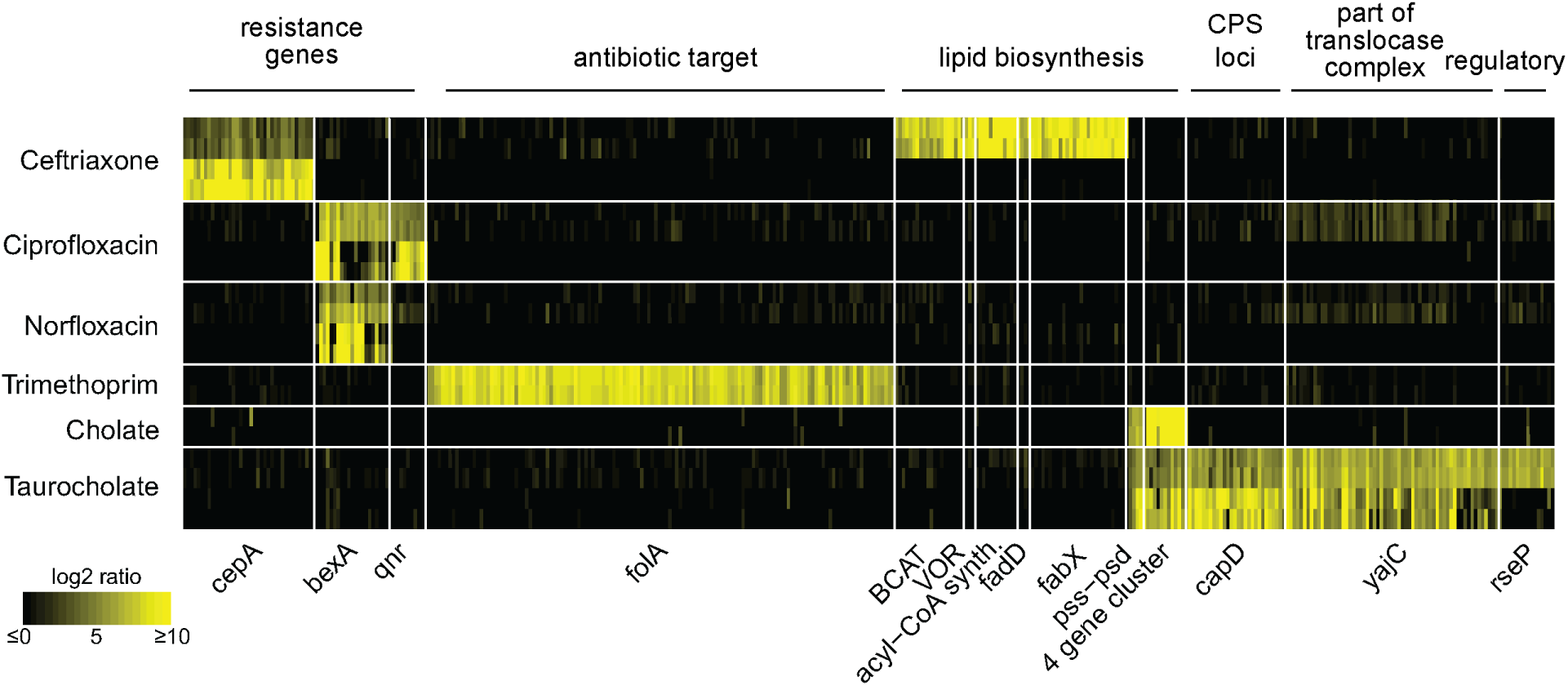
Selected gene hits that conferred benefit on antibiotics and bile salts. Fragments encoding these gene hits with significant high fitness scores from all Boba-seq libraries are displayed as fitness values. Replicates are plotted separately for each compound at either one or two concentrations. For compounds with hits at two concentrations, the lower concentration is plotted above the higher concentration. See Extended Data Table 5 for assay conditions, locus tags, and proposed mechanism of tolerance. Causative gene(s) across multiple libraries are plotted within their respective protein clusters.

Our data also associated five genes involved in the biosynthesis of branched-chain or unsaturated lipids with ceftriaxone tolerance, a third-generation cephalosporin in clinical use (Fig. 5, Extended Data Fig. 6). Collectively, a branched-chain amino acid aminotransferase (BCAT) and a 2-ketoisovalerate oxidoreductase (VOR) catalyze the conversion of branched-chain amino acids to ɑ-ketocarboxylic acids, which are oxidatively decarboxylated to form branched-chain acyl-CoAs. These can then be converted to acyl-ACPs and undergo elongation. In support of this, the BCAT homolog in *B. fragilis* (BF9343_3671, ∼88% identity) is involved in the biosynthesis of branched-chain ɑ-galactosylceramides, a type of immunomodulatory sphingolipids.^66^ We identified an acyl-CoA synthetase and a long-chain fatty acid-CoA ligase (FadD) in this condition as well (Extended Data Fig. 6, Fig. 6a). Specifically, biochemical characterization of a homolog of the acyl-CoA synthetase (Q8TLW1, ∼58% identity) revealed activity as a 2-methylbutyryl-CoA synthetase.^67^ Finally, a beneficial nitronate monooxygenase flavoprotein was identified to be FabX as it is similar to HP0773 (∼40% identity) from *Helicobacter pylori*.^68, 69^ FabX is essential for unsaturated fatty acyl biosynthesis and requires molecular oxygen as the electron acceptor *in vitro*.^68, 69^ However, a homolog in *Neisseria gonorrhoeae* (NGO1024, ∼36% identity) is essential for anaerobic unsaturated fatty acyl biosynthesis and relies on an unknown electron acceptor.^70^ We cloned out BCAT, VOR, and FabX and confirmed they are beneficial in individual growth assays (Fig. 6b, Extended Data Fig. 7d). Our results indicate that upregulating steps in branched-chain or unsaturated fatty acyl biosynthesis increases tolerance to ceftriaxone. Changes in the outer membrane lipid composition, including sphingolipids, is a potential strategy to reduce antibiotic concentrations in the periplasm where β-lactams act. However, the precise mechanism by which this is achieved warrants further work to decipher.

**Figure 6.**
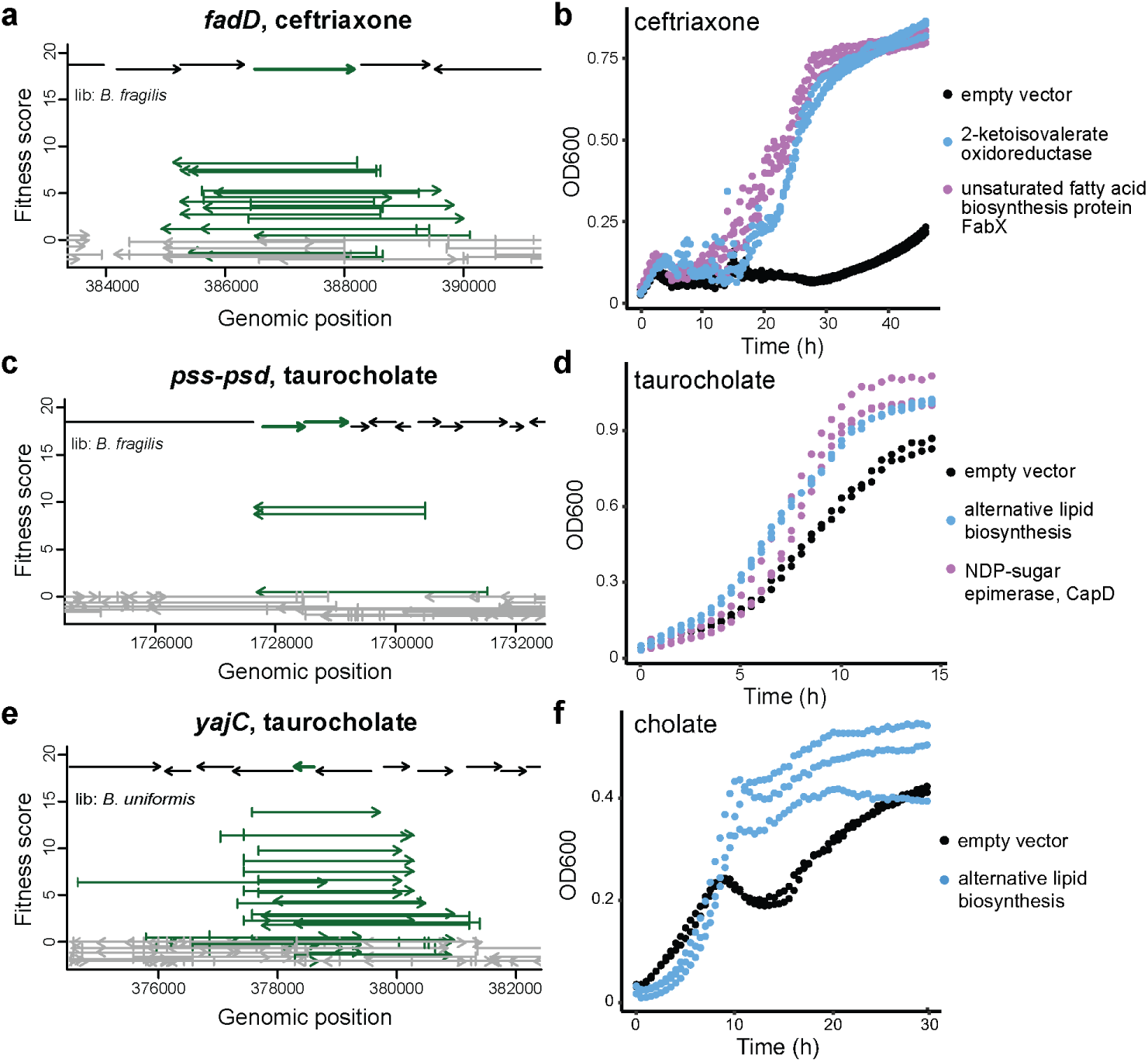
Genes involved in lipid biosynthesis, membrane protein translocation, and capsular polysaccharide biosynthesis increase tolerance to ceftriaxone, taurocholate, or cholate. Fragment plots for A) *fadD* (HMPREF1079_RS19625) in 0.5 mM ceftriaxone, C) phosphoethanolamine biosynthesis genes (HMPREF1079_RS14395, HMPREF1079_RS14390) in 10 mM taurocholate, and E) *yajC* (BACUNI_RS18600) in 10 mM taurocholate. Growth curves of individual hits expressed in *B. theta*: B) 2-ketoisovalerate oxidoreductase (HMPREF1077_RS19335, _RS17900, _RS17905, _RS17910) and *fabX* (HMPREF1077_RS10180) in 1 mM ceftriaxone; D, F) a gene cluster for alternative lipid biosynthesis (HMPREF1077_RS03620 to HMPREF1077_RS03635) in 5 mM cholate or 5 mM taurocholate; and D) a NDP-sugar epimerase *capD* (BACUNI_RS12910) in 5 mM taurocholate. All inserts include 200 bp upstream of the gene(s).

### Genes in lipid and surface polysaccharide biosynthesis contribute to bile salt tolerance

Bile salts are host-secreted factors that have antibacterial activities and can modulate host signaling pathways to drive dynamic changes in the gut microbiome. These molecules act as detergents to disrupt and destabilize membrane integrity. In the colon, bacteria may encounter bile salts up to 1 mM and encode various strategies to counter their toxicity.^71^ Here, we sought to identify factors that reduce susceptibility to bile stress in these microbes and thus may be important for colonization and competition in the humangut.

In both taurocholate and cholate, two beneficial loci are involved in lipid biosynthesis (Fig. 5). First, a two-gene cluster encoding phosphatidylserine synthase (Pss) and phosphatidylserine decarboxylase (Psd) catalyze the formation of phosphatidylserine and phosphatidylethanolamine (Fig. 6c, Extended Data Fig. 6). These genes are essential and cannot be studied using loss-of-function approaches, highlighting an advantage of using Boba-seq. Second, a conserved cluster of four genes (adenylyltransferase/cytidyltransferase, short-chain alcohol dehydrogenase/reductase, CDP-alcohol phosphatidyltransferase, lysophospholipid acyltransferase) is absent in *B. theta* and likely involved in the biosynthesis of an alternative lipid (Extended Data Fig. 6). This operon was enriched from the two *Parabacteroides* libraries and is prevalent among *Parabacteroides*, but not *Bacteroides*. We further confirmed its benefit for growth in taurocholate and cholate (Fig. 6d,f).

On the bile salt taurocholate, overexpression of RIP metallopeptidase RseP led to high fitness (Fig. 5). This is likely mediated through sigmaE factor activation by RseP in response to extracytoplasmic stress to induce transcriptional changes in cell surface proteins.^72^ Additional hits on taurocholate include the preprotein translocase subunit YajC and a NDP-sugar epimerase CapD (Fig. 5, 6e, Extended Data Fig. 6). YajC is a small inner membrane protein part of the translocase protein complex with SecDF, but it is not essential and its role is not well understood.^73^ In support of a role in responding to bile salts, protein levels of YajC in *Campylobacter jejuni* were elevated during exposure to chenodeoxycholate, a bile salt.^74^ NDP- sugar epimerase is often part of CPS loci in *B. theta*.^75^ The *cps1* locus in *B. theta* was induced in a rich medium by taurocholate but not by cholate^76^ and encodes a NDP-sugar epimerase (BT0381, >60% identity) similar to hits in our screen. Here, we further demonstrate that overexpressing the epimerase alone in *B. theta* is sufficient to increase tolerance to bile stress (Fig. 6d). Overall, both hits are supported by published gene expression data and might induce changes in membrane proteins or surface polysaccharides as a strategy to alleviate taurocholate toxicity.

In summary, our results connect genes involved in lipid biosynthesis, a regulatory pathway, membrane protein translocation, and surface polysaccharide biosynthesis to bile salt tolerance. While a few hits have supporting gene expression data from other studies, most genes are newly associated with this phenotype. Additionally, most studies quantify differentially abundant transcripts or proteins to uncover pathways involved in response to bile stress.^74, 76, 77^ Here, we leveraged high-throughput genetics to directly demonstrate that additional gene copies or the gain of a new gene cluster is sufficient to confer protection against bile salts.

## Discussion

Boba-seq is a new platform for constructing barcoded overexpression libraries for scalable and quantitative phenotypic screens. Here, we generated seven libraries using genomic DNA from *Bacteroides* and *Parabacteroides* species, an important clade of human gut bacteria. First, we validated our method in a complementation screen. Next, using wild-type *B. theta* as the expression host, we identified novel phenotypes in carbon utilization and stress tolerance across 120 unique regions and 22 compounds (Extended Data Table 5). By assaying DNA from multiple bacteria, we identified 30 hits supported by multiple protein homologs, which provided further confidence in our results.

Overall, we improved annotations and/or obtained new phenotypes for 29 protein clusters. In particular, we identified an outer membrane transporter for the uptake of raffinose, a plant-derived carbohydrate that is non-digestible by humans. This marks the first report of raffinose transporters in Bacteroidales. Using biochemistry and genetics, we identified a hexokinase critical for D-glucosamine metabolism. A region encoding a ssDNA-binding protein within a MGE led to a genomic recombination event and to a growth advantage on L-fucose, a key nutrient in the gut, although the exact mechanism remains unclear. Furthermore, we discovered genes that enhance tolerance toward antibiotics and bile salts, prevalent stressors encountered in the gut. Among these, genes involved in lipid biosynthesis, including a gene cluster absent in *B. theta*, increased tolerance to a cephalosporin and to bile salts. An epimerase that may alter surface polysaccharides was also beneficial in bile salts.

The selection of a physiologically related genetic background is critical for phenotypic assays. For instance, the kinase hits likely would not have been identified in an *E. coli* overexpression strain. *E. coli* uses the phosphotransferase system for D-glucosamine uptake where the substrate is phosphorylated during uptake, thus rendering the kinase activities redundant.^78^

We demonstrate the benefit of combining overexpression data with loss-of-function data to strengthen gene-to-phenotype mappings. 10 protein clusters have RB-TnSeq fitness data that are consistent with our Boba-seq data (Extended Data Table 5).^10, 11^ The scalability of both loss-of-function and gain-of-function libraries using barcoding techniques for Bacteroidales is an exciting development. By constructing RB-TnSeq and Boba-seq libraries for a large number of isolates and screening across identical conditions, we can rapidly generate specific functional hypotheses for novel genes.

A few variables can be adjusted in our workflow. Current libraries have fragment lengths of ∼3 kb and may not capture pathways with several proteins, therefore insert size can be increased to cover larger gene clusters. Additionally, the nature of competitive growth assays can lead to “jackpot” effects where the most fit strains outcompete other strains with a beneficial fragment and reduce the number of hits. This effect can be reduced by increasing the number of replicates, as BarSeq only costs a few dollars per sample, or by growing libraries on agar plates or in microfluidic droplets to spatially separate strains and prevent competition.^79, 80^

Boba-seq can be readily scaled to a larger number of libraries and assays. Our empty vector library has ∼14 million barcodes and although we multiplexed six libraries with ∼300,000 total strains, this can be increased. Libraries can be built using DNA from complex samples to access greater genetic diversity and uncultivated taxa. Moreover, our vectors replicate in at least nine additional Bacteroidales besides *B. theta*, which expands the diversity of functions that can be screened. Additionally, loss-of-function mutant strains can be used for high-throughput complementation screens. Finally, Boba-seq libraries can be tested in additional assay formats, including host colonization, phage infections, suppressors of toxic genes, and adhesion.^12, 25, 26, 53, 81, 82^

## Online Methods

### Strains and cultivation conditions

Strains, plasmids, and primers used in this study are listed in Extended Data Table 1. *E. coli* strains were routinely cultured in Luria-Bertani Lennox (LB) with the appropriate selection using 50 μg/mL carbenicillin (Carb50) or 50 μg/mL kanamycin (Kan50) at 37 °C with shaking at 200 rpm. *E. coli* WM3064 was grown with 300 μM diaminopimelic acid (DAP) supplement. Species within the order Bacteroidales were grown at 37 °C in an anaerobic chamber (Coy) maintained with a gas mix of 10% H_2_, 5% CO_2_, 85% N_2_. Media recipes and commercial sources of substrates are provided in Extended Data Table 4. Routine culturing was carried out in supplemented Brain Heart Infusion (BHIS) where BD Bacto Brain Heart Infusion was freshly supplemented with 0.5 mg/L hemin and 2.5 μL/L vitamin K_1_. The Varel-Bryant (VB) defined minimal medium was used for the majority of complementation and fitness assays as well as validation growth curves.^40^ B12 vitamin was excluded and instead, methionine was used in our medium. Cysteine was substituted with dithiothreitol (DTT) as the reducing agent for certain carbon conditions. An exception to this was the heparin validation growth curves, which were obtained in another defined minimal medium (MM).^44^ Media were made anaerobic by pre- reducing in the anaerobic chamber at least one day prior to use. Conjugation mixes were selected on 200 μg/mL gentamicin (Gm200) and 5 μg/mL erythromycin (Erm5) to obtain the correct mutant.

### Construction of replicative vectors for *B. theta*

The pNBU2_erm-TetR-P1T_DP-GH023 integrative vector was modified to generate replicative vectors. This vector contains a previously reported anhydrotetracycline (aTc) inducible promoter and RBS (P1T_DP-GH023).^33^ A PmlI restriction site, the synthetic terminator L3S2P21, and BsmBI sites used for barcoding were inserted downstream of the inducible promoter-RBS cassette.^29, 34, 35^ First, a BsmBI restriction site was removed from the pNBU2_erm-TetR-P1T_DP- GH023 vector using site-directed mutagenesis. The pNBU2 vector with no BsmBI site was linearized by PCR to exclude the attachment sites. Fragments containing replicative machinery and origin of replication from pTIO-1, pKSH24, and pFD340 were PCR amplified and purified using GFX DNA and Gel Band Purification kit.^31, 32, 83^ The third gene block consisted of an insertion site, a terminator, and a barcoding site.^10, 34, 35^ Three-part Gibson Assembly reactions were performed to generate pNBU2_repA1, pNBU2_repA2, and pNBU2_repA3 containing portions from pTIO-1, pKSH24, and pFD340, respectively. Gibson reactions were carried out according to manufacturer’s recommendations (NEB, Gibson Assembly master mix). This was transformed into electrocompetent *E. coli* EC100D *pir+* and plated onto LB-Carb50 to obtain the correct construct. All vectors were confirmed by Sanger sequencing.

### Detection of inducible expression using Luciferase assays

To quantify gene expression on the three pNBU2_repA vectors, luciferase assays were performed as previously reported, using Promega’s Nano-Glo Luciferase Assay System Kit.^33^ Briefly, the NanoLuc gene was inserted downstream of the aTc-inducible promoter via Gibson assembly. Sequence-verified vectors were conjugated into *B. theta*. These *B. theta* strains were inoculated into BHIS-Erm5 and grown anaerobically at 37 °C for 16–18 h. Starter cultures were subcultured into BHIS-Erm5 and varying concentrations of aTc at a 1:100 dilution to mid-exponential phase (OD of 0.4-0.6, 3–5 hours). 500 µL of each culture were then pelleted by centrifugation at 5,000 ✕ *g*, and cells were lysed with 50 µL of 1✕ BugBuster Protein Extraction Reagent (Sigma). Lysates clarified at 21,130 ✕ *g* for 5 minutes, and 10 µL of the supernatant was mixed with an equal volume of the NanoLuc reaction buffer containing the NanoLuc substrate. Luminescence was immediately measured using a BioTek Synergy H1 microplate reader (Agilent), and normalized to OD600 (optical density at 600 nm) of the culture at time of lysis.

### Determination of plasmid copy number using droplet digital PCR

Droplet digital PCR (ddPCR) was performed following a previously reported procedure to estimate the plasmid copy number of the pNBU2_repA vectors in *B. theta*.^84^ Briefly, *B. theta* containing each vector was grown in BHIS-Erm5 for 16–18 h at 37 °C, followed by a 1:100 dilution and subculture to mid-exponential phase (OD of 0.7–0.9, 3–5 h). 500 µL of each culture were then harvested for DNA extraction using Qiagen’s DNeasy blood & tissue kit, after which 1 µg of DNA was digested using NdeI (NEB). ddPCR was performed using approximately 0.01, 0.1, or 1 ng of the digested DNA as template for each 40 µL reaction. PCR was performed using the Evagreen supermix and primers p-AH204-p-AH207 (Extended Data Table 1), following BioRad’s protocols for multiplexing with Evagreen and using an annealing temperature of 60 °C. Samples were analyzed using Bio-Rad’s QX200 droplet digital PCR system. The resulting data points were manually gated, and reported concentrations for each amplicon were used in determining the plasmid copy number.

### Barcoding of the pNBU2_repA1 vector

Barcodes were amplified from synthesized DNA using Phusion polymerase (NEB) at an annealing temperature of 58 °C, elongation time of 60 s, and 6 cycles.^9^ pNBU2_repA1 vector was barcoded with random 20 nucleotides using Golden Gate Assembly. Each 20 μL reaction consisted of 1 μg of plasmid, 35 ng of barcode amplicons, 2 μL of T4 DNA ligase buffer, 1 μL of T4 DNA ligase (NEB), and 1 μL of BsmBI/Esp3I enzyme (Thermo). This resulted in a 2:1 molar ratio of insert to vector. Six reactions were incubated at 37 °C for 5 min and 16 °C for 10 min, which were repeated 10 cycles, followed by 37 °C for 30 min. All reactions were pooled, cleaned across two columns from the DNA clean and concentrator kit (Zymo), and eluted twice with 10 μL H_2_O. Uncut vectors were removed with additional incubations with BsmBI/Esp3I in 25 µL reactions according to the manufacturer’s protocol. Reactions were incubated at 37 °C for 2 h, after which 1 µL of BsmBI/Esp3I was added for incubation at 37 °C for 16 h. Each reaction was cleaned using DNA clean and concentrator kit (Zymo) and eluted twice with 8 μL H_2_O. 7 μL of cleaned reaction was transformed into 50 μL of electrocompetent *E. coli* WM3064 for a total of 4 transformations. All aliquots were recovered in 7 mL S.O.C. medium with 300 µM DAP for 1.5 h. An aliquot was removed for CFU counting and confirmed high transformation efficiency of 10^6–7^ CFUs per mL. Recovered cells were inoculated into 1 L of LB (50 μg/mL Carb, 300 μM DAP) for selection. Each vector library was purified for BarSeq to determine barcode diversity.

### Construction and quantification of barcoded genomic libraries

gDNA was purified from Bacteroidales using a DNeasy UltraClean Microbial Kit or DNeasy Blood & Tissue Kit (Qiagen). Up to 2 μg in 200 μL TE buffer was sheared by ultrasonication to an average of 3–5 kb with a Covaris S220 focused ultrasonicator using red miniTUBEs. Sheared gDNA was then gel-purified and size-selected by cutting out the 3–6 kb range. 4 µg of barcoded pNBU2_repA1 was digested in 50 µL reactions with 4.5 µL of PmlI. This was incubated at 37 °C for 30 minutes. Linearized vectors were dephosphorylated with 1 µL of rSAP (NEB) at 37 °C for 10 min and then deactivated at 65 °C for 5 min. Digested vectors were gel-purified using the Monarch DNA Gel extraction kit.

The *B. theta* library was constructed using blunt-end ligation, however vector rearrangement issues were observed by Sanger sequencing of individual *E. coli* clones. Cloning steps taken are similar to the previously reported Dub-seq library.^24^ Briefly, sheared and size-selected gDNA from *B. theta* was end-repaired using End-IT DNA End-Repair kit (Lucigen) in a 50 µL reaction consisting of 2 µL End-It Enz mix, 5 µL dNTP mix, 5 µL 10 mM ATP, 5 µL buffer, and 4 µg gDNA. This was incubated at room temperature for 45 min and deactivated at 70 °C for 20 min. This was gel-purified and size-selected for 3–6 kb fragments. The end-repaired, sheared DNA fragments were then ligated to linearized and dephosphorylated barcoded vectors. A 20 µL ligation reaction was performed with 10 µL Blunt/TA ligase master mix (NEB), 100 ng of vector backbone, and 4-fold molar excess of end-repaired DNA fragments, and incubated at room temperature for 20 min. Reaction was purified using Monarch PCR & DNA cleanup kit and eluted twice with 10 µL H_2_O. 20 ng of cleaned ligation reaction was transformed into 50 µL aliquots of commercial *E.coli* EC100D that was diluted 4-fold with 10% glycerol. A total of 8 transformations were performed and each recovered in 1 mL of S.O.C. medium at 37 °C for 1.5 h with shaking at 200 rpm. An aliquot was removed for CFU counting to estimate library diversity. Recovered cultures were combined and inoculated into 500 mL of LB-Carb50 for selection. The final library was purified for BarSeq to obtain an estimated diversity. The initial *B. theta* library consists of ∼440,000 barcodes and was diluted to obtain the final library with ∼70,000 barcodes.

An alternative strategy was applied to the remaining six libraries, which did not lead to any observable vector rearrangement. They were constructed by cloning DNA fragments into the pCR™Blunt II-TOPO™ vector using Zero Blunt™ TOPO™ PCR Cloning Kit (Invitrogen), followed by PCR amplification of the inserts and Gibson assembly into barcoded pNBU2_repA1. In this protocol, sheared gDNA is first end-repaired in 20 µL reactions using Quick CIP (NEB) and following the manufacturer’s protocol. End-repaired inserts are column-purified. Ligations were carried out in 6 µL TOPO reactions that contained 0.5 or 1 µL of pCR II-Blunt-TOPO vector, 1 µL of salt solution, and 20 ng of end-repaired inserts. Reactions were incubated at room temperature for 30 min and cleaned using SPRI beads (Beckman Coulter). Briefly, 0.9 equivalent volume of bead solution was mixed with each TOPO reaction and incubated for 15 min. Beads were washed three times with 80% EtOH and allowed to dry briefly. DNA was eluted with 5 µL of H_2_O heated to 50 °C and transformed into 50 µL of electrocompetent 3-fold diluted *E. coli* EPI300 (Lucigen). An aliquot was taken out for CFU counting on LB-Kan50. Recovered cultures were stored as glycerol stocks at −80 °C. Multiple TOPO reactions and transformations were carried out until an estimate of ∼40,000 to 100,000 CFUs was obtained for each library. Glycerol stocks of recovered cells from the same library were combined and inoculated into 200 mL of LB-Kan50 for selection. Inserts were PCR amplified from 150 ng of purified TOPO vectors using Q5 polymerase with an annealing temperature of 67 °C, 2 minute extension, and 8 cycles. Template vectors were digested using DpnI and the 3–6 kb size range was gel-purified. Insert amplicons were ligated into barcoded pNBU2_repA1 in 20 µL of Gibson reactions using the manufacturer’s protocol. 20 ng of cleaned reaction was transformed into 50 µL of 4-fold diluted electrocompetent *E. coli* EC100D. Cells were recovered and stored as glycerol stocks before selection. Aliquots of the recovered cells were plated onto LB-Carb50 to estimate CFUs. Libraries were serially diluted to <100,000 CFUs and selected in liquid LB-Carb50 to make final glycerol stocks and perform BarSeq. All libraries were transformed into *E. coli* WM3064 for conjugation into *B. theta*.

Differences between libraries constructed using each cloning workflow were observed. The ligated *B. theta* library has the highest percentage of genes covered (98%) and the lowest strand bias. The PCR required for Gibson Assembly, though with a low cycling number, introduced biases that likely led to lower genome coverage values (Table 1). Gibson-assembled libraries exhibited a slight strand bias toward fragments oriented opposite to the promoter (ratio of genes oriented opposite over genes oriented the same direction as vector promoter: 1.43– 1.88). In contrast, the number of genes on either strand is very similar in the *B. theta* library, with a ratio of 0.98. This is likely a result of gene toxicity in *E. coli* caused by the lac promoter on the TOPO vector. This strand bias did not increase upon library conjugation into *B. theta*, which is consistent with the low basal expression of the aTc inducible promoter.

### PacBio sequencing to map libraries

Barcoded libraries purified from EC100D were used as templates for amplicon sequencing. Each sample was dual indexed using 5’-phosphorylated PCR primers containing 16 nt barcodes.^85^ Barcode and insert regions were amplified in 50 µL PCRs with Q5 polymerase, 150 ng of vector template, 57 °C annealing temperature, 2 min extension time, and 6 or 8 cycles. 8– 10 reactions per library were pooled, digested with DpnI twice (NEB), and column-concentrated. Amplicons were size-selected for >800 bp lengths using 0.55 equivalent volume of AMPure PB beads (PacBio). 200–600 ng of this was used as sample input for the manufacturer’s protocol: Preparing SMRTbell libraries using PacBio Barcoded Overhang Adapters for Multiplexing Amplicons. The universal adapter was used during the ligation step instead of a barcoded adapter since amplicons are dual indexed. Adapter-ligated amplicons were digested with exonuclease using SMRTbell Enzyme Clean Up Kit 2.0 and purified using 0.55 equivalent volume of AMPure PB beads. Final amplicon solutions were quantified using Qubit BR Assay and sizes were confirmed on the Bioanalyzer HS DNA chip (Agilent). Samples were sequenced on the PacBio Sequel II instrument using SMRT cell 8M (QB3 Genomics, UC Berkeley). See Extended Data Table 2 for the number of demultiplexed CCS reads used to map each library. In general, a good run on one PacBio SMRT cell 8M was sufficient to map 3–4 libraries.

### Computational pipeline used to map barcodes to genes

The Boba-seq script used to map barcodes to genes and detailed explanations can be found at https://github.com/OGalOz/Boba-seq and as a part of the deposited data. A summary is provided as Extended Data Figure 4. Input data files are required and consist of: a reference assembly file (.fasta), a genome annotation file (.gff/.gff3), and long reads data (.fastq). The user also needs to provide a fasta file containing 4 short oligo sequences that flank the barcode and the insert fragment on each amplicon. A configuration .JSON file is used to set parameters for each step. Briefly, reads were first demultiplexed using PacBio’s lima tool. Barcode and insert sequences are extracted using 15bp flanking oligos using usearch (www.drive5.com/usearch/). Filters are applied to exclude reads with concatemers and incorrect barcode lengths. Inserts with an expected error of >10 are excluded using vsearch (https://github.com/torognes/vsearch). Inserts are mapped to the reference genome using minimap2 and results are kept for hits with high percent coverage and percent match values.^86^ The majority of barcodes mapped to a single genomic location. For barcodes that are mapped to multiple locations, most are due to sequencing errors and are within a few nt of each other. For barcodes mapping to overlapping locations, the location with the top read count is kept. Barcodes mapped to distant genomic locations are kept in the mapping tables, but are excluded from our fitness analysis. Finally, genomic positions are mapped to protein-coding genes to generate mapping tables, which are used to calculate library statistics.

### Conjugation into *B. theta* and other Bacteroidales

Single vectors were first transformed into conjugation donor strains *E. coli* S17-1 or WM3064. 40 mL of Bacteroidales recipient at OD ∼0.1 was mixed with 4 mL of *E. coli* donor at OD ∼1 outside of the anaerobic chamber. Mixed cultures were pelleted by centrifugation and resuspended in 250 μL BHIS. Each cell mixture was then pooled onto BHIS or BHIS-DAP (for WM3064) plates and incubated at 37 °C overnight aerobically for >16 h. The entire cell mass was then scraped and resuspended in 1 mL of BHIS, from which 20–100 µL was plated onto BHIS-Gm200-Erm5 agar. Antibiotic plates were incubated at 37 °C anaerobically for 48 h or longer until distinct colonies formed. Single colonies were restreaked onto BHIS-Gm200-Erm5 once, from which a single colony was then inoculated into BHIS-Gm200-Erm5. Bacteroidales cultures were inoculated into LB and incubated aerobically at 37 °C to ensure a lack of *E. coli* contamination. Glycerol stocks were checked by Sanger sequencing of plasmids and 16S rRNA. Using this protocol, pKSH24 and pTIO-1 were conjugated into *B. theta* to test replication in a new host. pNBU2_repA1, pNBU2_repA2, and pNBU2_repA3 with NanoLuc were conjugated into 13 different *Bacteroides*, *Parabacteroides*, and *Phocaeicola* species.

### Complementation libraries and assays

*B. theta, B. uniformis, B. fragilis*, and *P. johnsonii* libraries were conjugated into *B. theta* Δ*tdk*ΔBT2158 or Δ*tdk*ΔBT3703 using *E. coli* WM3064 as the conjugation strain. 200 µL of conjugation mix in 1 mL BHIS was inoculated into 100 mL of pre-reduced BHIS-Gm200-Erm5 in the anaerobic chamber. An aliquot of the conjugation mix was serially diluted for CFU counting to ensure that ∼10^6^ transconjugants were inoculated for selection. Cultures at OD >1 were pelleted and resuspended in BHIS with 0.15% Cys and 15% glycerol to make glycerol stocks.

For complementation assays, the starter culture of each library was diluted to an OD of 0.5 in BHIS-Gm200-Erm5 with 40 ng/mL aTc and incubated at 37 °C for 3.5 h to induce gene expression. Cells were pelleted and washed with VB minimal medium twice; these served as Time0 samples. Each library was inoculated at a starting OD of 0.02 in 1 mL VB-Erm5-aTc40 and 20 mM of either leucrose, palatinose, trehalose, or ɑ-cyclodextrin. Cultures were incubated at 37 °C and harvested when turbid. Each assay was also streaked onto BHIS-Gm200-Erm5 agar plates. Colony PCR and Sanger sequencing were used to identify clones containing BT2158, BT4448, or BACUNI_RS05440.

### Growth assays of complemented mutants

Mutants complemented with BT2158, BT4448, or BACUNI_RS05440 (Extended Data Table 6), *B. theta* Δ*tdk*ΔBT2158, and *B. theta* Δ*tdk* parental strain were inoculated into BHIS-Erm5 or BHIS and grew overnight at 37 °C. Cells were pelleted and washed once with VB minimal medium. Each mutant was inoculated at a starting OD of 0.05 in 200 µL VB-Erm5 and 20 mM trehalose. *B. theta* Δ*tdk* and *B. theta* Δ*tdk*ΔBT2158 were inoculated in the same manner, except without Erm. Growth measurements were obtained in a 96-well plate using an Epoch 2 Microplate Spectrophotometer (Agilent) inside of an anaerobic chamber. OD600 values shown on plots are pathlength-corrected and blank-normalized.

### Fitness screen of pooled libraries in wild-type *B. theta*

*B. caccae, B. fragilis*, *B. salyersiae, B. uniformis, P. johnsonii,* and *P. merdae* libraries were conjugated into wild-type *B. theta* as described for complementation libraries. For fitness assays, 1 mL of glycerol stock was pelleted and resuspended in 10 mL of BHIS-Erm5 with 40 ng/mL of aTc as starter cultures to grow for 16–18 h. These were then pelleted and washed with VB-Erm5 twice. Libraries were pooled equally based on OD to a final OD of 5, which served as Time0 samples. For carbon utilization assays, pooled libraries were inoculated at a starting OD of 0.04 in 1 mL of VB-Erm5 with 40 ng/mL aTc, and a single carbon substrate at the concentration listed in Extended Data Table 4. For stress tolerance assays, pooled libraries were inoculated at a starting OD of 0.1 or 0.04 in 0.5 mL of BHIS-Erm5, 40 ng/mL aTc, and an inhibitory compound at a range of concentrations listed on Extended Data Table 4. Assays were carried out in 48-well plates and incubated at 37 °C. Each condition was set up in replicates of two on the same day. A few conditions were repeated on a second day of experimentation where new conditions were tested as well. Cultures were harvested when turbid for BarSeq.

### DNA barcode sequencing (BarSeq) and analysis

Vector libraries were purified from *E. coli* using E.Z.N.A. Plasmid DNA Mini Kit (Omega Bio-tek) or Monarch Plasmid Miniprep kit (NEB). However, this led to low yields and high levels of contaminants when extracting from Bacteroidales. DNA purified from the same set of Bacteroidales samples using plasmid purification kits and genomic DNA purification kits were tested for barcode PCR amplification. BarSeq results from both purified plasmids and gDNA were comparable and therefore, DNeasy Blood & Tissue Kit or QiaAmp DNA QIAcube HT Kit (Qiagen) for 96-well sample plates were used to extract DNA from *B. theta* cultures for BarSeq for all fitness assays.

Barcodes were amplified using the previous-published “BarSeq_V3” primers or using new “BarSeq_V4” primers (Extended Data Table 1).^9, 87^ The BarSeq version 4 primers are similar to the BarSeq version 3 primers but use a 10-bp index sequence for P7 primers and a 8-bp index for P5 (instead of 6 bp for both in version 3) and allow multiplexing up to 768 samples (instead of 96). Briefly, BarSeq PCRs were performed as 50 µL reactions with Q5 polymerase (NEB) and 200 ng of template DNA: 98 °C for 4 min, 25 cycles of 98 °C for 30 s, 55 °C for 30 s, and 72 °C for 30 s, and 72 °C for 5 min. For both BarSeq3 and BarSeq4 designs, the P7 (or P2) BarSeq primers allowed demultiplexing by Illumina software,^9^ and the P5 (or P1) BarSeq primers contained an additional index checked by the MultiCodes.pl script.^87^ All PCRs were pooled equally in volume and column-cleaned. Up to 96 samples (BarSeq3) or 152 samples (BarSeq4) were multiplexed for sequencing on full lanes of Illumina HiSeq400 SE50 (QB3 Genomics, UC Berkeley), partial lanes of NovaSeq6000 PE150 (Novogene), or full lanes of MiSeq (v2 reagent kit). 11,000–94,000 reads per sample were obtained for complementation assays sequenced on the MiSeq and 0.3–5.3 million reads were obtained for samples sequenced on the HiSeq or NovaSeq.

BarSeq reads were converted to counts per barcode using the MultiCodes.pl script in the feba code base (https://bitbucket.org/berkeleylab/feba/), with the -minQuality 0 option and either -bs3 or -bs4 for each version of BarSeq primers. To estimate the diversity of barcoded libraries, only reads that had a quality score of ≥30 at each position (the -minQuality 30 option) were used, which corresponds to an error rate for barcodes of at most 0.001 * 20 nt = 2%. Furthermore, any barcodes that were off-by-1 errors from a more common barcode were eliminated. Barcodes observed in the PacBio dataset are compared to barcode sequences from the BarSeq data to determine the percentage of barcodes mapped.

For the highly diverse empty library, the Chao1 estimator was used to estimate the barcode diversity from 2.99 million high-quality reads,^88^ The Chao1 estimator uses the number of barcodes seen just once, but some may be due to sequencing errors. To correct for this, barcodes that were off-by-1 nucleotide from a more abundant barcode were ignored, and the number of singletons was reduced by an assumed error rate ranging from 0% to 2%. This yielded estimates ranging from 13.5–14.1 million barcodes for the empty pNBU2_repA1 library.

### Calculation of fitness scores for each fragment

Only barcodes that mapped to a single location in one library and that were present in the conjugation donor (*E. coli* WM3064) libraries were considered. 2% of barcodes mapped to more than one library and another 5% of mapped barcodes were not detected in the BarSeq data for the conjugation donor libraries. For complementation assays, we computed fitness scores for all of these fragments. For fitness assays in the wild-type *B. theta* background with six pooled libraries, 280,036 of the 305,382 barcodes (that were present in WM3064) were detected in the Time0 samples. Also, barcodes that were not detected in Time0 samples occasionally had statistically significant benefits. This is likely due to positive selection that led mutants to gain in abundance in the assay condition despite their low initial abundance. Therefore, fitness scores were computed for all mapped barcodes that were present in WM3064. The script used to calculate fitness scores can be found at https://github.com/morgannprice/BobaseqFitness.

The fitness of each strain was defined as:

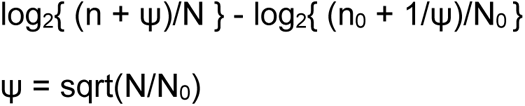

where n is the number of reads for this barcode in the experimental sample, n_0_ is the number of reads for this barcode across the Time0 samples, and N and N_0_ are the total number of reads for all considered barcodes in the experimental or Time0 samples, respectively. The pseudocount ψ is added to the counts to ensure that the log ratio is not undefined if either count is zero. This value of ψ also ensures that if both counts (n and n_0_) are 0, then the fitness will be zero (corresponding to no evidence of a change in relative abundance). If the experimental and Time0 samples have the same total number of reads, then ψ = 1.

### Determination of statistically significant hits

The minimum fitness value threshold for statistical significance was selected based on a control comparison between the two Time0 samples collected on the same day for the large fitness screen. Each sample contained a mixture of six genomic libraries and had 2.9 million or 3.0 million total reads. Of the 305,382 barcodes considered, the fitness values ranged from −3.9 to +4.1. Therefore, a minimum fitness value of +5 was selected for a statistically-significant hit or barcode.

Because a few complementation assays were sequenced at much lower coverage, a z score cutoff was also applied. The standard error of the fitness value was estimated as the square root of the variance given Poisson noise in counts^9^, or

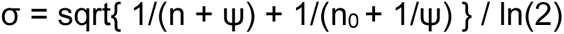

The z score is then fitness / σ and z ≥ 4 is selected as the cutoff. For the standard normal distribution and 305,382 barcodes, 10 values would be expected above this threshold.

### Selection of biologically consistent hits and the likely causative protein for each region

We focused on barcodes and inserts that had significantly high fitness scores in both replicate experiments conducted in the same condition and on the same day. We considered these barcodes to have a biologically consistent phenotype if either an overlapping fragment or a fragment encoding a similar protein had statistically significant benefits. Specifically, an insert was confirmed by overlap if another insert from the same region of the genome was significant in both replicates. An insert was confirmed by similarity if the likely causative protein for the region (see below) was similar (at least 40% identity and 75% coverage) to a protein that was found within another insert that had significantly high fitness in two replicates for the same condition. For experiments in the wild-type *B. theta* background where libraries were pooled, the confirmation would occur in the same experiments. For complementation assays where libraries were assayed separately, the criterion might be fulfilled from different assays in the same condition.

For each region with inserts that were statistically significant in both replicates, the insert with the highest fitness score (after averaging replicates) was selected first. For each protein encoded within this insert, we computed several scores. First, we counted the number of significantly-beneficial inserts that contain this protein (with each replicate counted separately). Second, we counted the number of similar proteins that were in high-scoring regions. Third, we computed the average fitness score for all inserts containing this gene. To select the likely causative protein for each region, we chose the one with the highest number of beneficial inserts, or the highest number of similar proteins, or the highest average fitness. We used the additional scores to break ties.

For each of the putative causative proteins, we visually inspected the plot of fitness scores versus insert fragments. When available, we also considered the plots for similar experiments (such as the same carbon source with a different base medium, or a different concentration of the stress, or the same condition conducted on a different day) with similar protein hits. In 20 cases, we determined that more than one protein is necessary to confer a strong benefit. In 6 cases, we identified a different protein as conferring the benefit. In 28 cases, we were not sure which protein(s) conferred the benefit and we left these hits uncurated in Extended Data Table 5. For the remaining 162 cases, the automatically-selected protein was deemed correct.

### Validation of gene hits using individual strains

Selected gene hits from the fitness screens were PCR amplified from the corresponding gDNA, cloned into the pNBU2_repA1 backbone via Gibson Assembly, and transformed into *E. coli* S17-1 to generate validation vectors. Mutants with high fitness scores were selected for validations and the exact fragment boundaries for these were extracted from library mapping tables to design primers (Extended Data Table 6). Additionally, amplicons of the precise gene or genes were obtained. Validation vectors were conjugated into *B. theta*, and each strain was grown in the same condition as carried out for the fitness assay with AraC hits being an exception. Here, we switched to a different defined minimal medium (MM) where *B. theta* grew more quickly in heparin (see Extended Data Table 4 for composition).^44^ Each strain was cultured in 200 µL on a 96-well plate in replicates of 3. In a few cases, the empty vector control was grown up in replicates of 2. Growth measurements were obtained using an Epoch 2 Microplate Spectrophotometer (Agilent). OD values are pathlength-corrected and blank- normalized.

### Overexpression and purification of hexokinase

Hexokinase (INE72_RS21610) was amplified from *B. caccae* CL03T12C61 and cloned into pET29b vector for protein overexpression and purification. 500 mL of *E. coli* BL21(DE3) transformed with pET29b-hexokinase was induced with 0.5 mM IPTG at OD ∼0.6. Pellet was harvested and then resuspended in lysis buffer (50 mM Tris HCl pH 7, 100 mM NaCl, 1 mM PMSF, 0.2 mg/ml lysozyme, 1X SIGMAFAST protease inhibitor, 1% streptomycin sulfate, 5 mM BME). Resuspended cells were lysed using a sonicator probe. The clarified supernatant was loaded onto equilibrated Ni Sepharose 6 Fast Flow resin (Cytiva) and washed with a buffer containing 50 mM Tris HCl pH 7, 100 mM NaCl, and 5 mM BME. Protein was eluted using the wash buffer with 10, 20, 50, 100, or 200 mM imidazole and fractions were collected at each concentration. All fractions were visualized on SDS-PAGE and the appropriate fractions were pooled and concentrated down using 30K MWCO concentrators. Concentrated protein eluate was desalted using a PD-10 column loaded with Sephadex G-25 resin (Cytiva). Protein concentrations were calculated using Abs280 measurements from Nanodrop and a molar extinction coefficient of 44,350 M^−1^ cm^−1^. In total, 4.2 mg of protein was purified to a concentration of 105 μM. Protein aliquots were stored at –80°C.

### Hexokinase enzyme assays

*In vitro* coupled enzyme assays were performed to test substrate preference of hexokinase. Assay components include 50 mM Tris-Cl pH 7, 50 mM NaCl, 2 mM MgSO_4_, 1 mM NADH, 2 mM phospho(enol)pyruvic acid tri(cyclohexylammonium) salt (PEP), 1 mM ATP, and 1 mM substrate. Glucose, mannose, D-glucosamine, and N-acetyl- D-glucosamine were tested as substrates. Enzymes were added to concentrations of 10 U/mL lactate dehydrogenase from rabbit muscle (L2500-10KU, Sigma), 10 U/mL pyruvate kinase from rabbit muscle (P1506-1KU, Sigma), and 5 μM purified hexokinase. Lactate dehydrogenase and pyruvate kinase concentrations were also tested at a 2-fold increase to ensure they were in excess and not the rate limiting step. Assays were initiated by adding hexokinase and incubated for 5 minutes at room temperature. Assays were performed in volumes of 40 uL in triplicates in a 384-well plate. NADH consumption was observed through pathlength-corrected absorbance at 340 nm measured using a BioTek Synergy HTX plate reader.

### RNA-seq analysis of *ssb* region in L-fucose

For RNA-seq experiments, a *B. theta* strain expressing the *ssb* fragment 2 (Extended Data Table 6) and a strain containing the empty vector were grown in replicates of 3 in BHIS-Erm5- aTc40 to mid-exponential phase (OD600 = 0.4–0.8) and harvested using RNAprotect bacterial agent (Qiagen) and the RNeasy Mini kit (Qiagen) combined with on-column DNA depletion using the RNase-free DNase set (Qiagen). ∼500 ng of total RNA from each sample was submitted to Novogene for further processing; this included rRNA depletion followed by library preparation with the NEBNext Ultra II Directional RNA Library Prep Kit for Illumina (NEB). Library quality was checked using a 2100 Bioanalyzer (Agilent).

Sequencing reads were trimmed and aligned to the *B. theta* VPI-5482 genome (GCF_000011065.1) using HISAT2 (2.2.1) with the strand-specific option for paired-end reads, and default parameters otherwise.^89^ Mapped reads were then assigned to genome features using featureCounts with the strand specific setting.^90^ Due to concern over repeat regions, counts were generated with and without multimapping using featureCounts, which led to similar results. Final counts were generated with multimapping. These counts were normalized and analyzed using the DESeq2 package in R, resulting in log2 fold-changes along with test statistics and adjusted *p*-values generated using the Wald test (Extended Data Table 7).^91^

### Materials and data availability

The script for mapping Boba-seq libraries is available at https://github.com/OGalOz/Boba-seq. The script for calculating fitness scores is available at https://github.com/morgannprice/BobaseqFitness. Plasmid maps, a snapshot of the python script used for library mapping, mapping tables for each library, a snapshot of the R script used for fitness score calculations along with the input files, output files from fitness analysis, and raw reads from the RNA-seq experiment are deposited at: https://doi.org/10.6084/m9.figshare.24195054. All other data are available from the authors upon request. Please direct plasmids and strain requests to Y.Y.H. and A.P.A.

## Acknowledgements

We thank Dr. Naoya Ohara (Okayama University) for pTIO-1, Dr. Andrei Shkoporov (University of College Cork) for pKSH24, and Dr. Laurie Comstock (University of Chicago) for pFD340. pNBU2_erm-TetR-P1T_DP-GH023 and pNBU2_erm-TetR-P1T_DP-GH023-NanoLuc were gifts from Dr. Andrew Goodman at Yale University (Addgene plasmids #90324 and #117728). We thank Surya Tripathi for access to unpublished RB-TnSeq fitness data, and Bradley Biggs, Shekhar Mishra, Vivek Mutalik, and Valentine Trotter for helpful feedback on the manuscript. Y.Y.H. is an Astellas Pharmaceuticals Awardee of the Life Sciences Research Foundation, which funded part of this work. A.H. is an awardee of NSF GRFP grant DGE-2146752. We are grateful for the support of the Dean A. Richard Newton Memorial Professor Chair Funds. This work was supported in part by ENIGMA – Ecosystems and Networks Integrated with Genes and Molecular Assemblies (http://enigma.lbl.gov), a Science Focus Area Program at Lawrence Berkeley National Laboratory, supported by the U.S. Department of Energy, Office of Science, Office of Biological & Environmental Research under contract number DE-AC02-05CH11231. This work was partly supported by the NIH grant RM1 GM135102 (to A.M.D.). We acknowledge QB3 Genomics, UC Berkeley, Berkeley, CA for sequencing support through RRID:SCR_022170. This work was supported by NIH S10 OD018174 Instrumentation Grant. Several Bacteroidales used in this study were obtained through BEI Resources, NIAID, NIH as part of the Human Microbiome Project.

## Author contributions

Y.Y.H., M.N.P., A.M.D., and A.P.A. conceived the project. Y.Y.H., A.H., A.M.D., D.H., and H.C. carried out experimental work and collected data. Y.Y.H., M.N.P., A.H., and O.G. performed computational analyses. Y.Y.H. and M.N.P. wrote the original draft. Y.Y.H., M.N.P., A.H., A.M.D., and A.P.A. revised and edited the paper.

## Extended Data Figures

**Extended Data Figure 1.**
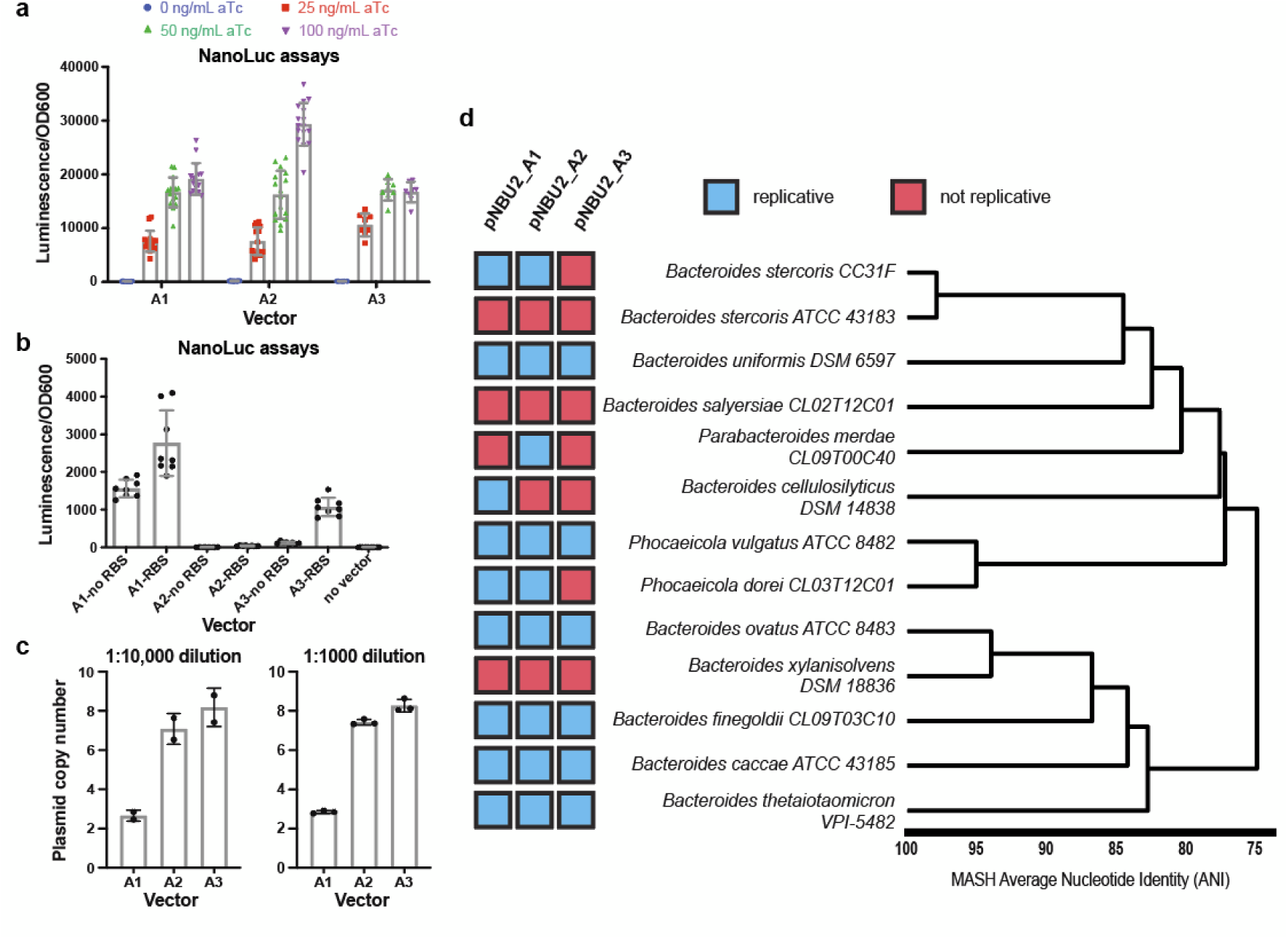
Anhydrotetracycline (aTc) induces expression of NanoLuc on pNBU2 replicative vectors in B. theta. A) A range of aTc concentrations (0, 25, 50, 100 ng/mL) was tested for three vectors with NanoLuc inserted downstream of the inducible promoter and an RBS. A total of two or four experiments were performed per vector with each experiment consisting of four replicates. B) NanoLuc was inserted into the pNBU2_repA1 vector in the opposite orientation to the aTc inducible promoter to test for constitutive promoter activity downstream of the insert site. NanoLuc gene with and without the upstream RBS (GH023) were tested in the absence of aTc. Two experiments were carried out with each experiment consisting of four replicates. Luminescence values were normalized by OD600 for all assays. Standard deviation is shown as error bars for both plots. C) Plasmid copy numbers in B. theta determined using droplet digital PCR (ddPCR). Cultures in the late exponential phase were harvested for this assay. Replicates from two dilutions were used to calculate plasmid copy numbers. D) Host range of pNBU2-based vectors across 13 Bacteroidales. Genome distance estimation and visualization were performed using dRep.^92^

**Extended Data Figure 2.**
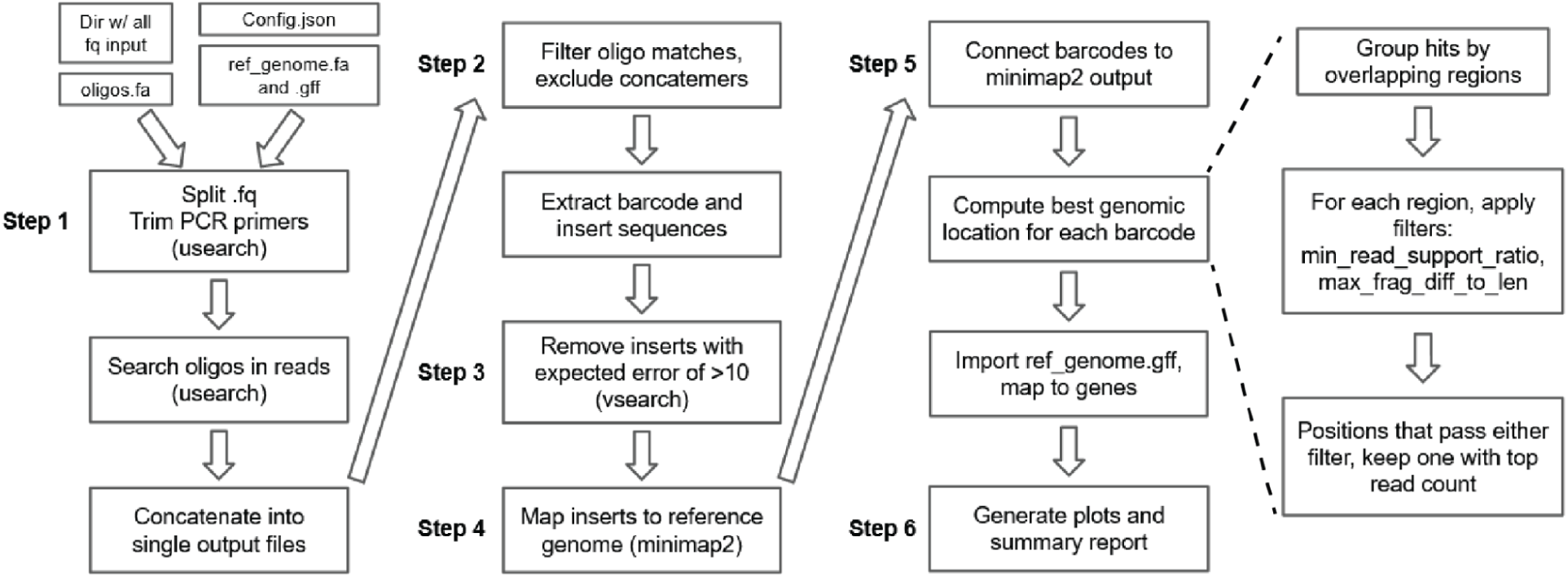
Overview of the Boba-seq mapping script. Input files, tools, and all steps are listed. Additional details for step 5 are listed for barcodes that map to multiple, genomic regions. Although the empty vector library has a high barcode diversity, it is possible to clone multiple inserts into vectors sharing the same barcode. We observed this in a very small fraction of barcodes and later removed them from our fitness score analysis, but they are reported by the mapping script. Two parameters are used to compute the best genomic location for barcodes that map to very similar positions. First, the ratio of highest read count to the second highest read count (min_BC_support_ratio value) is set at a threshold of 3 to identify the position supported by more reads. Second, the max_frag_diff_to_len value is the fraction of the maximum distance between start and end positions of 5’ of mapped locations plus the maximum distance between start and end positions of 3’ of mapped locations within each region, over the average insert length. This is set to a threshold of 0.25 in our current pipeline. All filters and parameters can be readily adjusted in the configuration .JSON input file.

**Extended Data Figure 3.**
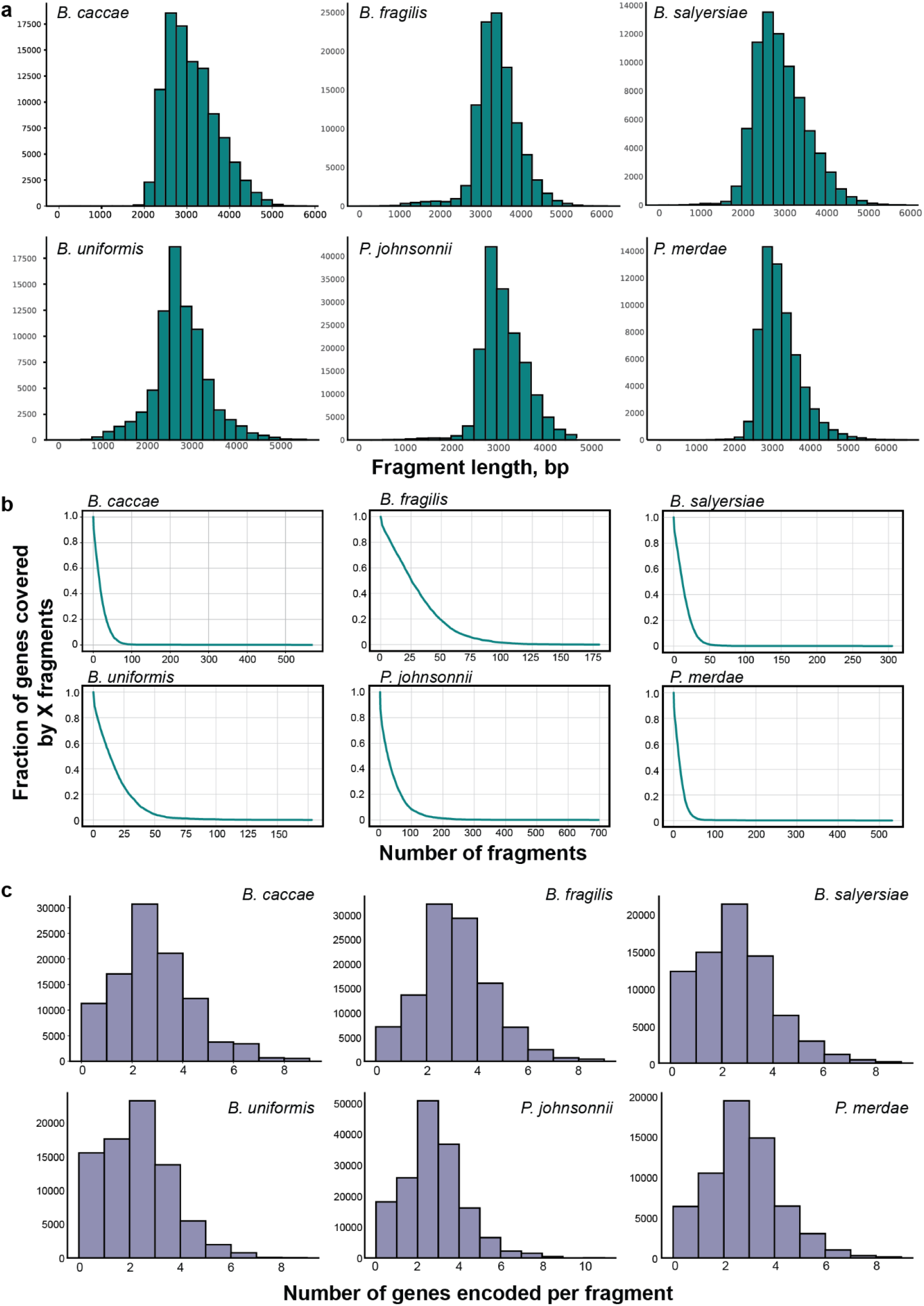
Summary plots for six Boba-seq genomic libraries. Plots are generated as part of the mapping script. A) Histograms of fragment sizes. B) Cumulative gene coverage plots. C) Histograms of the number of full-length genes encoded per fragment.

**Extended Data Figure 4.**
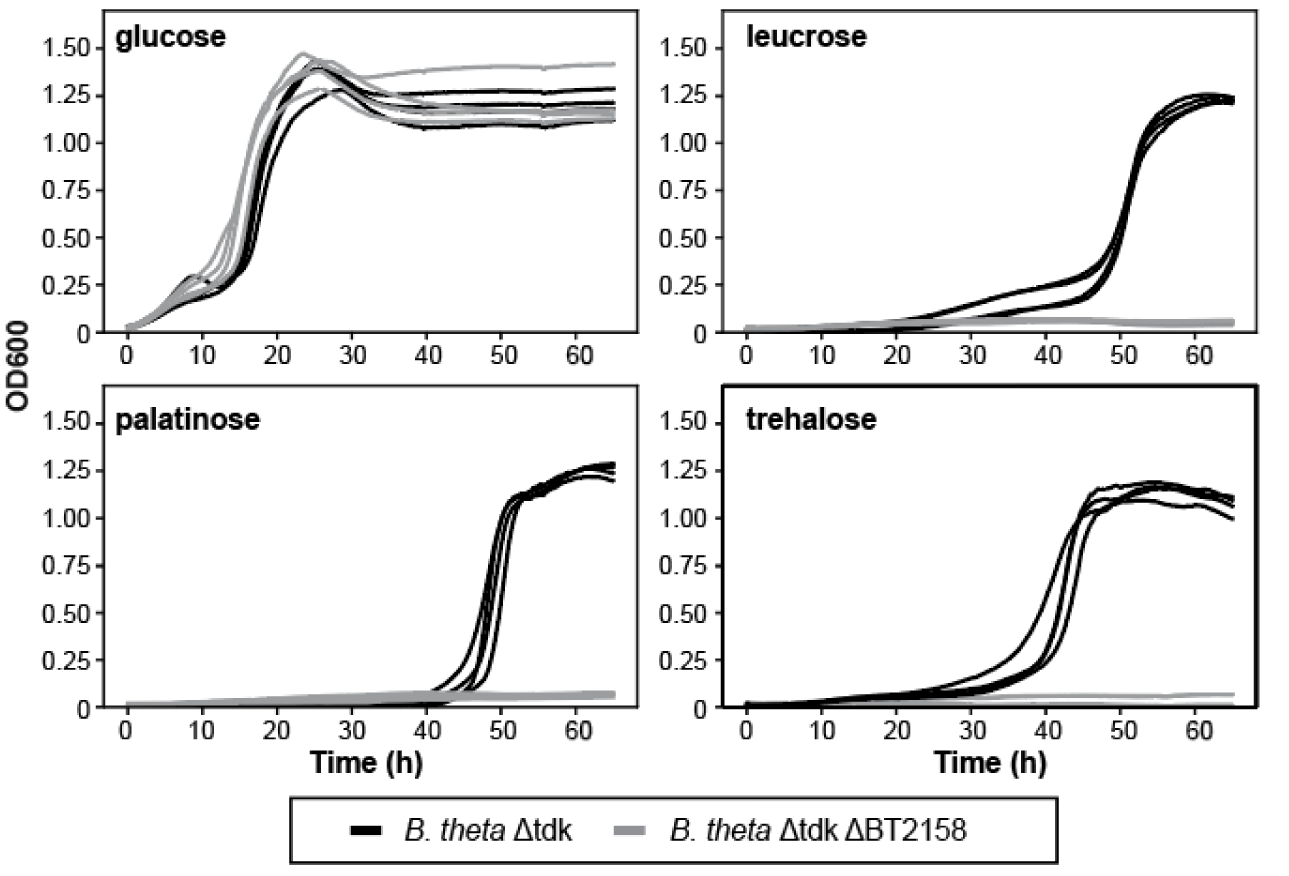
Growth curves of *B. theta* Δ*tdk* parental strain and Δ*tdk*ΔBT2158 mutant on 20 mM glucose, leucrose, palatinose, or trehalose in the VB minimal medium. Each strain was grown in replicates of 4 per condition.

**Extended Data Figure 5.**
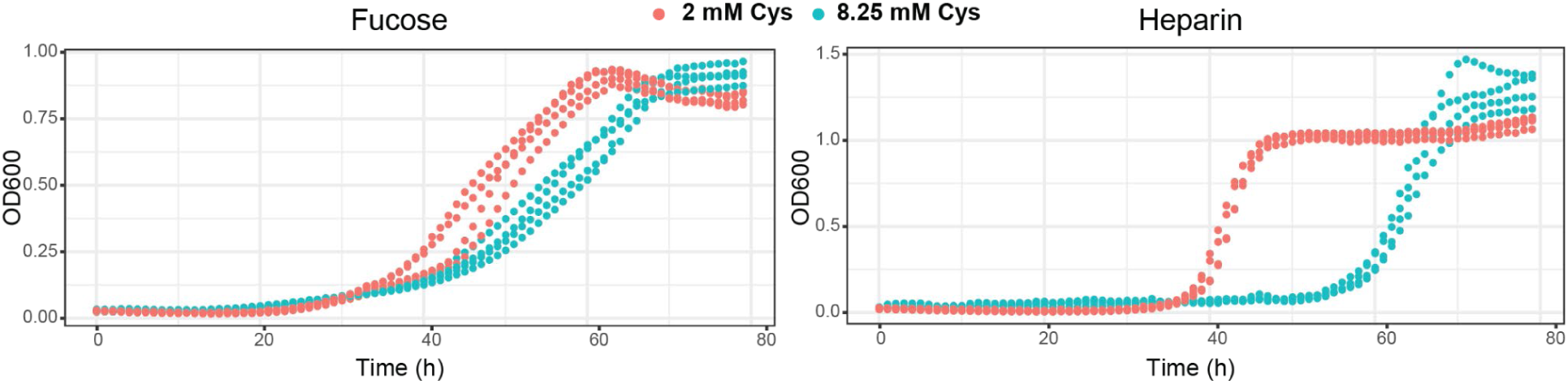
Growth of wild-type *B. theta* is inhibited by the high cysteine concentration (8.25 mM) in the VB minimal medium. Examples are shown for 20 mM L-fucose and 10 mg/mL heparin in replicates of 4. A lower cysteine concentration (2 mM) resulted in improved growth.

**Extended Data Figure 6.**
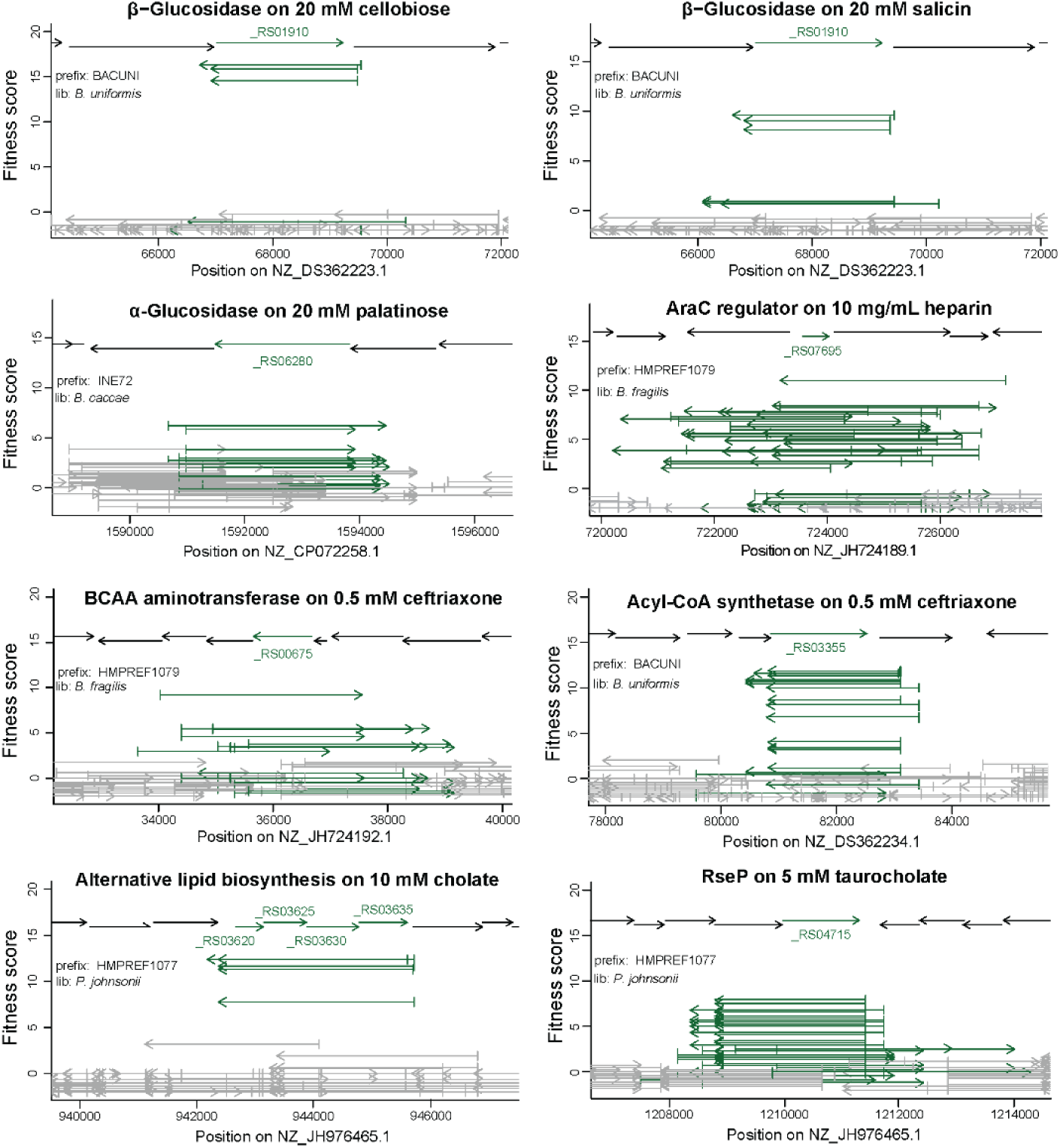
Fitness versus fragment plots of gene hits across carbon utilization and stress assays. Each arrow represents a mapped fragment in the corresponding Boba-seq library. The beneficial gene(s) and all inserts covering the full-length gene(s) are highlighted in green. The genomic positions of the inserts within the source genome are displayed on the x-axis. The average fitness score of each fragment from two experiments are shown.

**Extended Data Figure 7.**
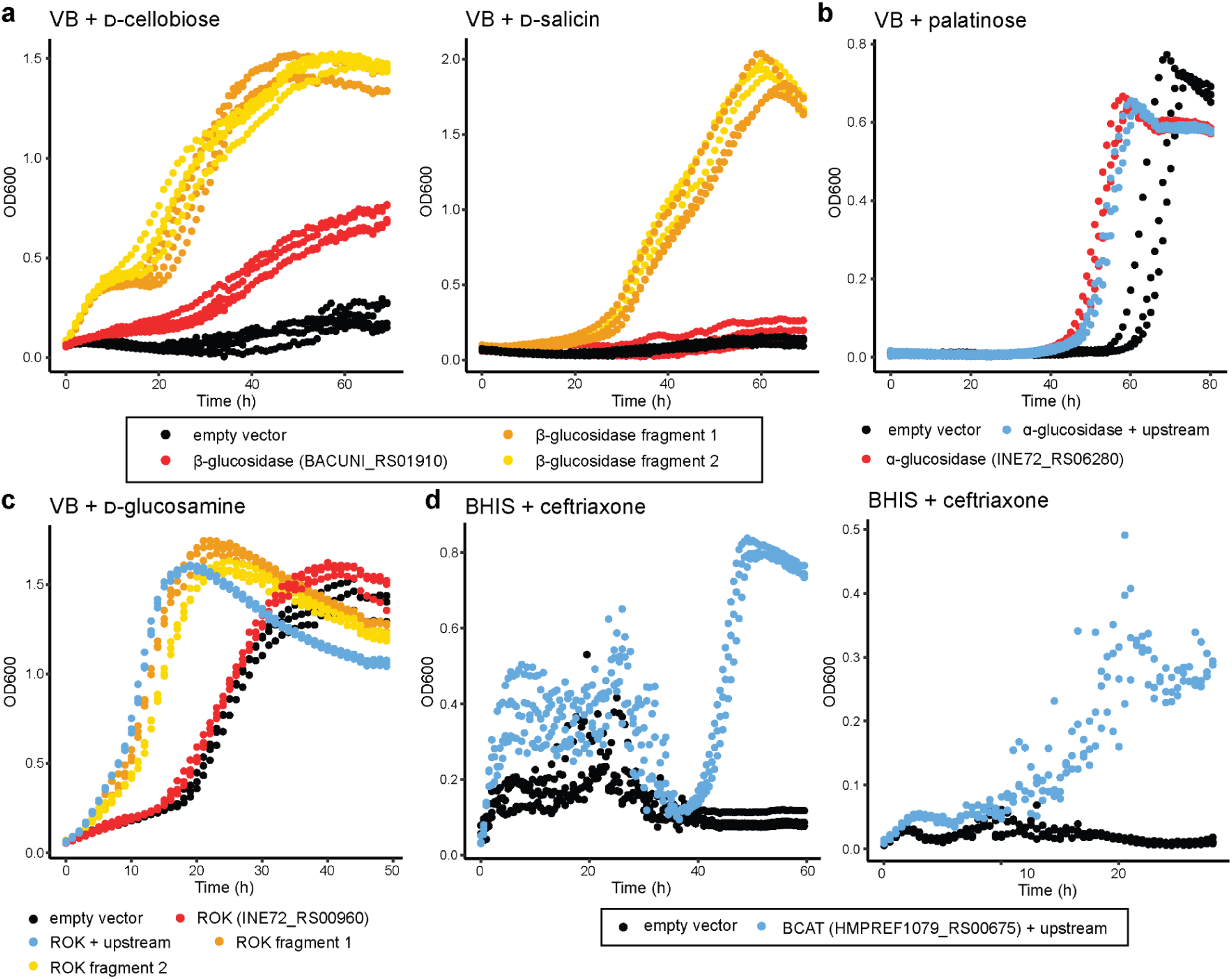
Growth benefits conferred by gene hits in individual *B. theta* strains. Each strain is conjugated with either a vector encoding the causative gene, the causative gene and the upstream 200 bp (or 100 bp for BCAT), a complete insert fragment from the library, or an empty vector. See Extended Data Table 6 for precise genomic boundaries, orientations, and fitness scores for each library fragment. Strains were constructed to confirm hits from A) D-cellobiose and D-salicin, B) palatinose, and C) D-glucosamine carbon substrates (20 mM) in the VB defined minimal medium. D) A branched-chain amino acid aminotransferase gene was confirmed to provide a benefit in BHIS with 1 mM ceftriaxone in two experiments. OD values are pathlength-corrected and blank-normalized. Each strain was grown in replicates of 3.

**Extended Data Figure 8.**
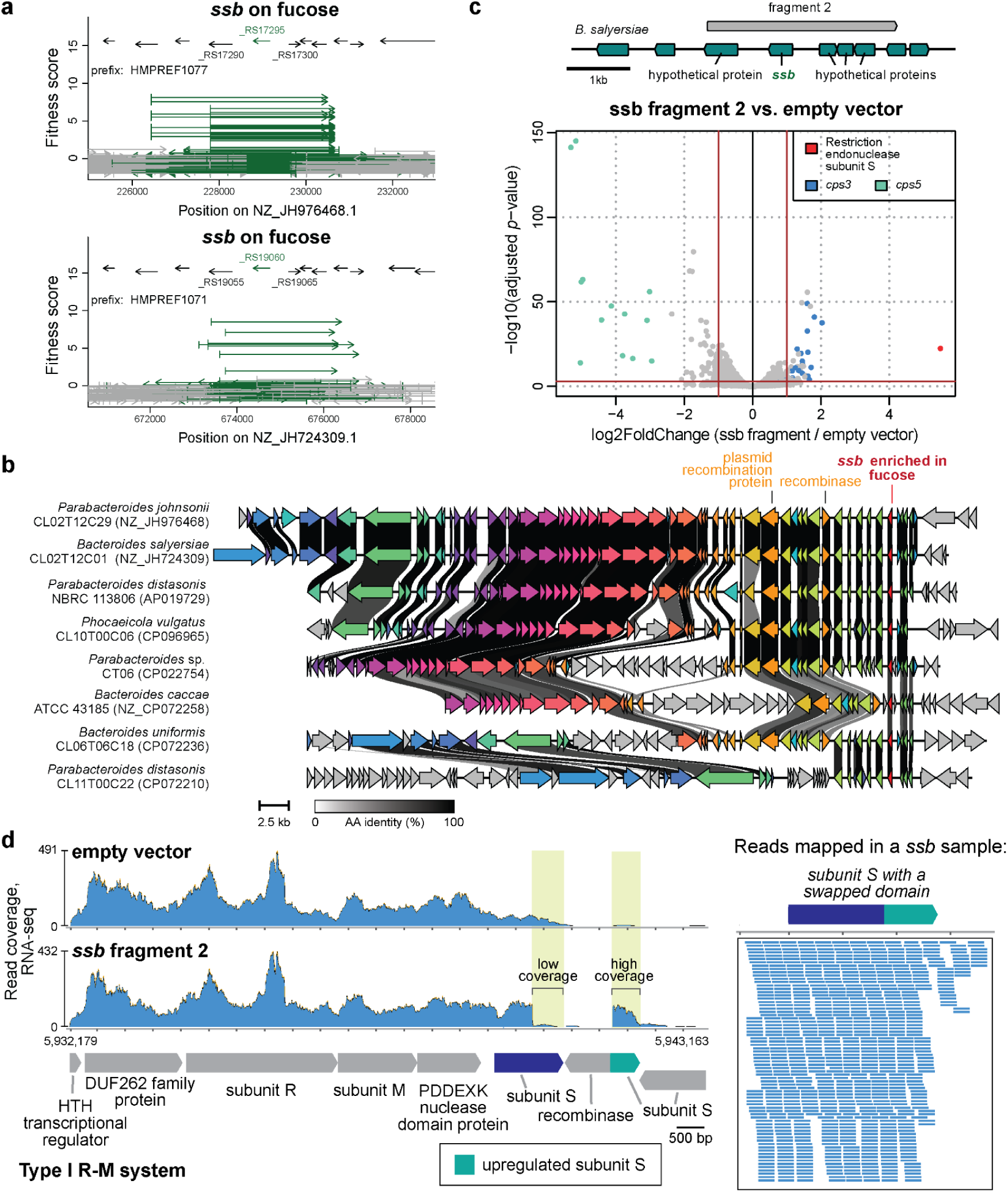
A genomic region in two different isolates (*B. salyersiae* and *P. johnsonii*) encoding an SSB protein conferred a strong benefit for growth on 20 mM L-fucose. A) The fitness versus fragment plots for both regions. The fitness scores displayed are averages across 8 experiments, including assays with different reductants (Cys or DTT) and in the presence or absence of aTc. B) Gene neighborhoods of the *ssb* region among Bacteroidales suggest this region is part of a mobile genetic element inserted at different genomic locations. Insertions and rearrangements are observed downstream of the *ssb* gene along with a conserved plasmid recombination protein and a recombinase. Genomic comparisons were visualized using Clinker with an amino acid identity cutoff of 40%.^93^ C) Log2 ratios of fold-change and adjusted *p*-values (Wald test) from the RNA-seq experiment of *B. theta* encoding the ssb fragment 2 compared to an empty vector. Selected hits are labeled by color. D) Coverage plots of sense strand reads from RNA-seq data point to a genomic rearrangement that led to the upregulation of a subunit S (BT_RS22805/BT4522) part of a type I R-M system. Alignments were made using HISAT2 with default splice parameters on the strand-specific setting. Representative plots were generated using sense strand reads from empty_1 and ssb_1 samples. Sense strand reads from ssb_1 sample (*B. theta* encoding the *ssb* fragment 2) map across the subunit S gene formed after recombination.

